# Does insulin signalling decide glucose levels in the fasting steady state?

**DOI:** 10.1101/553016

**Authors:** Manawa Diwekar-Joshi, Milind Watve

## Abstract

Recent work has suggested that altered insulin signalling may not be central to the pathophysiology of type 2 diabetes as classically believed. We critically re-examine the role of insulin in glucose homeostasis using five different approaches (i) systematic review and meta-analysis of tissue specific insulin receptor knock-outs, (ii) systematic review and meta-analysis of insulin suppression and insulin enhancement experiments, (iii) differentiating steady-state and post-meal state glucose levels in streptozotocin treated rats in primary experiments (iv) mathematical and theoretical considerations and (v) glucose insulin relationship in human epidemiological data. All the approaches converge on the inference that although insulin action is hastens the return to a steady-state after a glucose load, there is no evidence that insulin action determines the steady-state level of glucose. The inability to differentiate steady state causality from perturbed state causality has led to misinterpretation of the evidence for the role of insulin in glucose regulation.

## 1. Introduction

### Why is insulin believed to regulate fasting blood sugar: A burden of history?

After the classical demonstration by Claud Bernard that damage to medulla oblongata causes hyperglycaemia (1), the second major breakthrough was the demonstration by von Mering and Minkowski that pancreatectomy resulted in hyperglycaemia (2) and further that pancreatic extracts resulted in lowering of plasma glucose. The active principle eventually purified became known as insulin (3). The discovery and success of insulin in treating diabetes was so overwhelming that insulin became the key molecule in glucose homeostasis and the role of brain and other mechanisms were practically forgotten. It should be noted that the prevalent type of diabetes then was what we would call type 1 diabetes (T1D) today in which there is almost complete destruction of pancreatic β-cells. The distinction between type 1 and 2 developed gradually over the next five decades along with the realization that insulin levels may be normal or raised in type 2 diabetes (T2D) and that a substantial population of β-cells survives lifelong (4–6). However, by now the thinking about glucose homeostasis was so insulin centred, that the inability of normal or raised levels of insulin to keep plasma glucose normal was labelled as “insulin resistance” without adequately examining and eliminating alternative possibilities and the concept got wide uncritical acceptance. Although insulin resistance as a phenomenon is well established and its molecular mechanisms elucidated with substantial details, the question whether altered insulin signalling is solely or mainly responsible for fasting hyperglycaemia of T2D, or other insulin independent mechanisms play a significant role is not clearly answered.

There are multiple reasons to doubt and re-examine the role of insulin in glucose regulation in relation to T2D (7–9). Exogenous insulin and other insulin-centred lines of treatment have largely failed to reduce diabetic complications and mortality in T2D although short term glucose lowering may be achieved (10–15). In the long run even the glucose normalization goal is not achieved in majority of cases (12, 14). A number of mechanisms are known to influence glucose dynamics, partially or completely independent of insulin signalling, including autonomic signals (16, 17), glucocorticoids (18–21), insulin independent glucose transporters (22) and certain other hormones and growth factors (23–26). Analysis of multi-organ signalling network models have also raised doubts about the central role of insulin and insulin resistance in T2D (27).

The definitions as well as clinical measures of insulin resistance are such that the effects of all other mechanisms are accounted for under the name of “insulin resistance”. For example, the HOMA-IR index is calculated as a product of fasting glucose and fasting insulin (28, 29). The belief that this product reflects insulin resistance is necessarily based on the assumption that insulin signalling alone quantitatively determines glucose level in a fasting steady state. The assumption has seldom been critically examined. If any other mechanisms are contributing to raised fasting glucose levels, they will be included in the HOMA-IR index going by the way it is calculated and would be labelled as insulin resistance. We have previously showed using mathematical and statistical tools of causal analysis (30) that the classical pathway of obesity induced insulin resistance leading to a hyperinsulinemic normoglycemic prediabetic state and the faithfulness of HOMA indices in measuring insulin resistance cannot be simultaneously true. Either the HOMA indices do not represent insulin resistance faithfully or the classically believed pathway of compensatory insulin response leading to hyperinsulinemic normoglycemic state is wrong according to this analysis (30).

We examine here the long held belief that altered insulin signalling is responsible for fasting as well as post prandial hyperglycemia in T2D using five different approaches (i) Systematic review and meta-analysis of experiments involving tissue specific insulin receptor knock-outs (IRKOs) (ii) Systematic review and meta-analysis of experiments to chronically raise or lower insulin levels (iii) Primary experiments on streptozotocin (STZ) induced hyperglycaemia in rats that differentiate between steady and perturbed-state (iv) Examining the insulin resistance hypothesis for being mathematically possible and theoretically sound (v) Analysis of insulin-glucose relationship in steady-state versus post-meal perturbed-state in human epidemiological data for testing the predictions of mathematical models. The first three approaches have the advantage of using specific molecular interventions where the target is precisely known. For the meta-analyses we chose mechanisms of insulin level/action modification which have been used extensively and have been reproduced by multiple labs world over. The possible disadvantage is that they are mostly animal experiments and doubts are expressed about whether the results are directly relevant to humans (31–33). However, some of the experiments reported are human and they converge with the inferences of the animal experiments. In the last two approaches, human epidemiological data are used in which the experimental molecular precision is not expected, but we test certain specific predictions of the insulin resistance hypotheses using novel analytical approaches and examine whether they converge on similar inferences. The convergence of human and animal data is important to reach robust conclusions.

### 2. Systematic review and meta-analysis of experiments involving tissue specific insulin receptor knock-outs

The first step in insulin signalling is the binding of insulin to insulin receptor (34). The downstream actions of this event finally lead to insulin-dependent glucose uptake in insulin dependent tissues of the body. Experimentally, disruption of insulin signalling is achieved by knocking out or inhibiting various players in the signalling cascade. We chose to look at the effects of knocking out the tissue specific insulin receptor on fasting and post-meal or post glucose load levels in rodent models. Studying tissue specific insulin receptor knockouts enables us to differentiate between the roles of insulin signalling in different tissues. A classical belief is that the post-meal glucose curve is mainly influenced by the rate of glucose uptake by tissues, mainly muscle, whereas the fasting glucose levels are mainly determined by the rate of liver glucose production (35). If this belief is true one expects that muscle specific knockout would mainly affect the GTT curve but may not affect fasting glucose level, whereas liver specific knockout would mainly affect the fasting glucose level.

### 2.1. Methods

The details of the systematic literature review are given in table 1. The details of the experiments of the shortlisted studies can be seen in the table 1 of the supplementary information 1 which shows that similar methods have been utilised to create the knockouts and therefore a comparative analysis is justified.

**Table 1:** Systematic literature review for studies on tissue specific insulin receptor knockouts (meta-analysis 1).

| Meta-analysis 1→<br>Task performed ↓ | Insulin receptor knockout |
| --- | --- |
| Key word(s) used for the first search on the PubMed/MEDLINE data-base | “insulin receptor knockout” |
| Number of hits in the first search | 78 |
| Inclusion criteria for primary screening | Study showing experiments with IRKOs in which GTT curve has been reported |
| Number of papers shortlisted based on primary screening | 36 |
| Inclusion criteria for secondary screening | Study showing similar methods of making the insulin receptor knockout; had fasting and post glucose bolus readings of the control and knockout |
| Number of papers shortlisted based on screening the full-text and back referencing (data for meta-analysis extracted from these papers) | 16 |
| List of publications used in the final meta-analysis: references numbers | (36–51) |

#### 2.1.1 Statistical approach for meta-analysis

Although we short-listed papers that used similar methods, small differences in protocols can make considerable differences. As the results will reveal, there is substantial variation in results across studies. Therefore we use non-parametric methods for analysing the pooled data. We first look at in how many of the experiments the treatment mean in greater than the control mean and in how many it is less. If this difference is significant, we conclude that there is sufficient qualitative consistency across experiments to reach a reliable inference. If there is a consistent direction of difference, we look at how many are individually significant. As a conservative approach we avoid pooling data quantitatively since across studies there are differences in age or weight of animals, number of days after treatment, number of hours of fasting and other variables. This approach is maintained throughout all meta-analyses reported.

### 2.2 Results of the IRKO meta-analysis

We shortlisted 16 papers with 46 independent experiments in which glucose tolerance curves of insulin receptor knockouts and controls were compared (table 1 of supplementary information 1). The experiments could be segregated in four different tissue specific knockouts for the analysis: Liver insulin receptor knockout (LIRKO), Muscle insulin receptor knockout (MIRKO), fat/adipose insulin receptor knockout (FIRKO) and β-cell insulin receptor knockout (βIRKO). A generalized trend in the total picture summed up over all four IRKOs seen in the meta-analysis was that along the GTT curve, significantly higher glucose levels are seen in the knockouts as compared to the controls, particularly and consistently at 30, 60 and 120 minutes. However, the fasting glucose level was not significantly different in the meta-analysis. In some studies, fasting glucose was significantly greater in the knockouts than the controls, however in some other studies it is significantly lower as well. In 29 out of 46 experiments there was no significant difference (Table 2) in the fasting glucose levels of knockouts and controls. This trend was consistently seen in MIRKO, LIRKO and βIRKO. Only in FIRKO there were greater number of studies showing fasting glucose significantly higher in the knockouts than in the controls, but in the non-parametric meta-analysis the trend was not significant. Also, only in FIRKO, the 30, 60- and 120-minute glucose was not significantly different in the knockouts than the controls. It is notable in particular that in none of LIRKO experiments the fasting sugar was significantly higher than the controls. This contradicts the classical belief that liver insulin resistance is mainly responsible for fasting hyperglycaemia in T2D (35, 52).

**Table 2.** Meta-analysis of the fasting and post-feeding glucose levels in the control and IRKOs. The table shows, out of the total number of experiments used for the analysis, in how many the mean of the knockouts (T) was greater than the control means (C) and in how many the trend was reverse. This relative position of the means across studies is compared non-parametrically to see whether the trend across studies was non-random, significant ones being indicated by asterisk. The table also gives in how many studies T was significantly greater than C and vice versa. It can be seen that for fasting glucose the difference is not significant in majority of studies and where there is statistical significance there is lack of consistency across studies. However, at 30, 60 and 120 minutes the knockouts have consistently elevated levels of glucose as compared to the corresponding controls.

|  | **Total studies** | **T>C** | **T<C** | ***p* using chi square** | **T > C individually significant** | **T < C individually significant** |
| --- | --- | --- | --- | --- | --- | --- |
| **All IRKOs** |  |  |  |  |  |  |
| Fasting | 46 | 25 | 20 | 0.454 | 13 | 4 |
| 15 minutes | 14 | 7 | 7 | 0.999 | 4 | 2 |
| 30 minutes | 40 | 36 | 4 | <0.0001* | 22 | 1 |
| 60 minutes | 40 | 36 | 4 | <0.0001* | 24 | 1 |
| 120 minutes | 46 | 37 | 9 | <0.0001* | 24 | 2 |
| **FIRKO** |  |  |  |  |  |  |
| Fasting | 12 | 9 | 3 | 0.083 | 9 | 1 |
| 15 minutes | 3 | 1 | 2 | 0.566 | 1 | 1 |
| 30 minutes | 9 | 7 | 2 | 0.095 | 6 | 1 |
| 60 minutes | 9 | 7 | 2 | 0.095 | 5 | 1 |
| 120 minutes | 12 | 9 | 3 | 0.83 | 7 | 2 |
| **MIRKO** |  |  |  |  |  |  |
| Fasting | 10 | 3 | 7 | 0.205 | 0 | 2 |
| 15 minutes | 6 | 3 | 3 | 0.999 | 1 | 0 |
| 30 minutes | 10 | 9 | 1 | 0.011* | 3 | 0 |
| 60 minutes | 10 | 9 | 1 | 0.011* | 3 | 0 |
| 120 minutes | 10 | 6 | 4 | 0.527 | 3 | 0 |
| **LIRKO** |  |  |  |  |  |  |
| Fasting | 9 | 4 | 5 | 0.739 | 0 | 0 |
| 15 minutes | 1 | 1 | 0 | N.A. | 0 | 0 |
| 30 minutes | 9 | 9 | 0 | 0.003* | 6 | 0 |
| 60 minutes | 9 | 9 | 0 | 0.003* | 7 | 0 |
| 90 minutes | 9 | 9 | 0 | 0.003* | 5 | 0 |
| 120 minutes | 9 | 7 | 2 | 0.094 | 4 | 0 |
| **βIRKO** |  |  |  |  |  |  |
| Fasting | 8 | 6 | 2 | 0.157 | 4 | 0 |
| 15 minutes | 2 | 2 | 0 | 0.157 | 2 | 0 |
| 30 minutes | 8 | 7 | 1 | 0.033* | 6 | 0 |
| 60 minutes | 8 | 7 | 1 | 0.033* | 7 | 0 |
| 90 minutes | 4 | 4 | 0 | 0.046* | 4 | 0 |
| 120 minutes | 8 | 8 | 0 | 0.0046* | 7 | 0 |

A possible problem in comparing fasting glucose across different studies was that different fasting intervals have been used ranging from 4 to 16 hours. No study clearly reported how much time is required to reach a steady-state in a knockout. In 10 of the experiments in which fasting time was reported as 16 hours, none had fasting sugar significantly different for controls. In the 13 experiments in which it was high, the fasting duration was between 4 to 12 hours or not precisely reported. Therefore, it is likely that in some of the experiments, glucose steady-state was not yet achieved at the time point defined as fasting. This bias increases the probability that higher fasting glucose is reported for the knockouts. However, since we do not see a significant difference in the meta-analysis, the inference that IRKO does not alter fasting glucose is unlikely to be a result of the bias. In fact, any possible correction to the bias might further reduce the apparent residual difference. Therefore, in spite of some inconsistency across studies, a robust generalization is that IRKOs have significantly increased plasma glucose over controls at 30 to 120 minutes post glucose load but they do not appear to affect steady-state fasting glucose. The time required to reach the steady state is nevertheless increased.

## ***3.*** Systematic review and meta-analysis of insulin increase and insulin suppression experiments

The insulin receptor knockout experiments are based on the assumption that the main action of insulin is through the specific receptors. It can be argued that insulin acts through other receptors or may have other mechanisms of action yet unknown and therefore receptor knockouts do not fully eliminate insulin action. Alternatively, we can alter insulin level itself to see how it affects glucose level in fasting state or post glucose load. Insulin is known to alter plasma glucose immediately on administration but this is not a steady-state response. If insulin levels can be raised or lowered and sustained long enough to reach a steady state, the effect of insulin on glucose in a steady-state can be studied. If insulin affects steady-state glucose, a sustained rise in insulin will result into a sustained lower steady-state glucose level. Conversely a sustained suppression of insulin would lead to higher steady-state glucose. We studied published literature for experiments where a stable and sustained increase or decrease in insulin was achieved and then the effect on fasting glucose and GTT studied.

### ***3.1*** Methods

#### 3.1.1 Increase in insulin

The model of choice for a sustained increase in insulin levels is a knock out or inhibition of the insulin degrading enzyme (IDE). An interplay between insulin secretion and insulin degradation maintains the level of insulin in plasma (53–56). Plasma insulin has a half-life of 4 to 9 minutes (57, 58) and it is degraded predominantly by the insulin degrading enzyme (IDE) (54, 57). Inhibition of IDE has been considered as a therapeutic option for type 2 diabetes with limited success (59, 60). We performed a systematic literature review to find out experiments in which IDE was inhibited to obtain a sustained high plasma insulin level and, in such animals, GTT was performed (table 3).

**Table 3:**
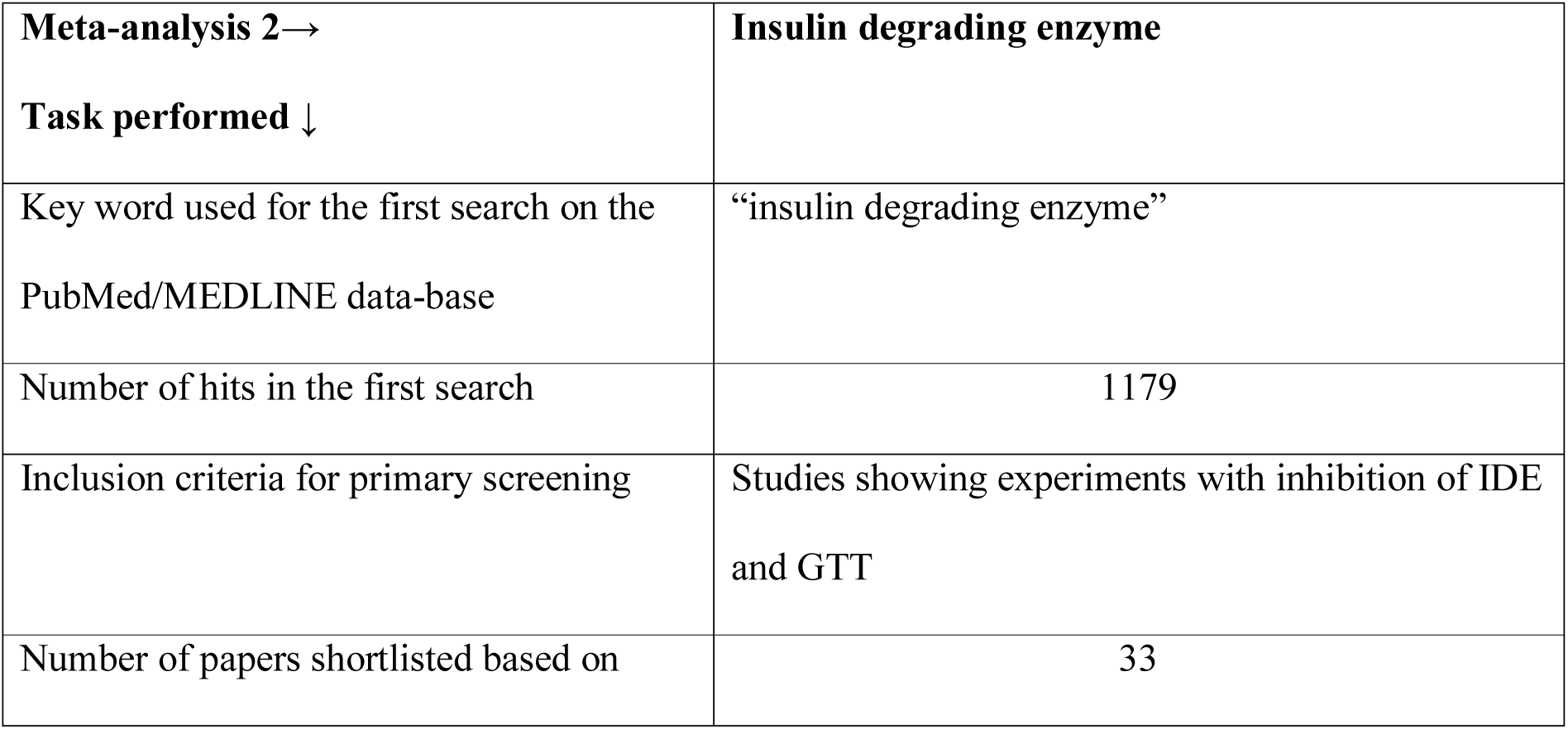
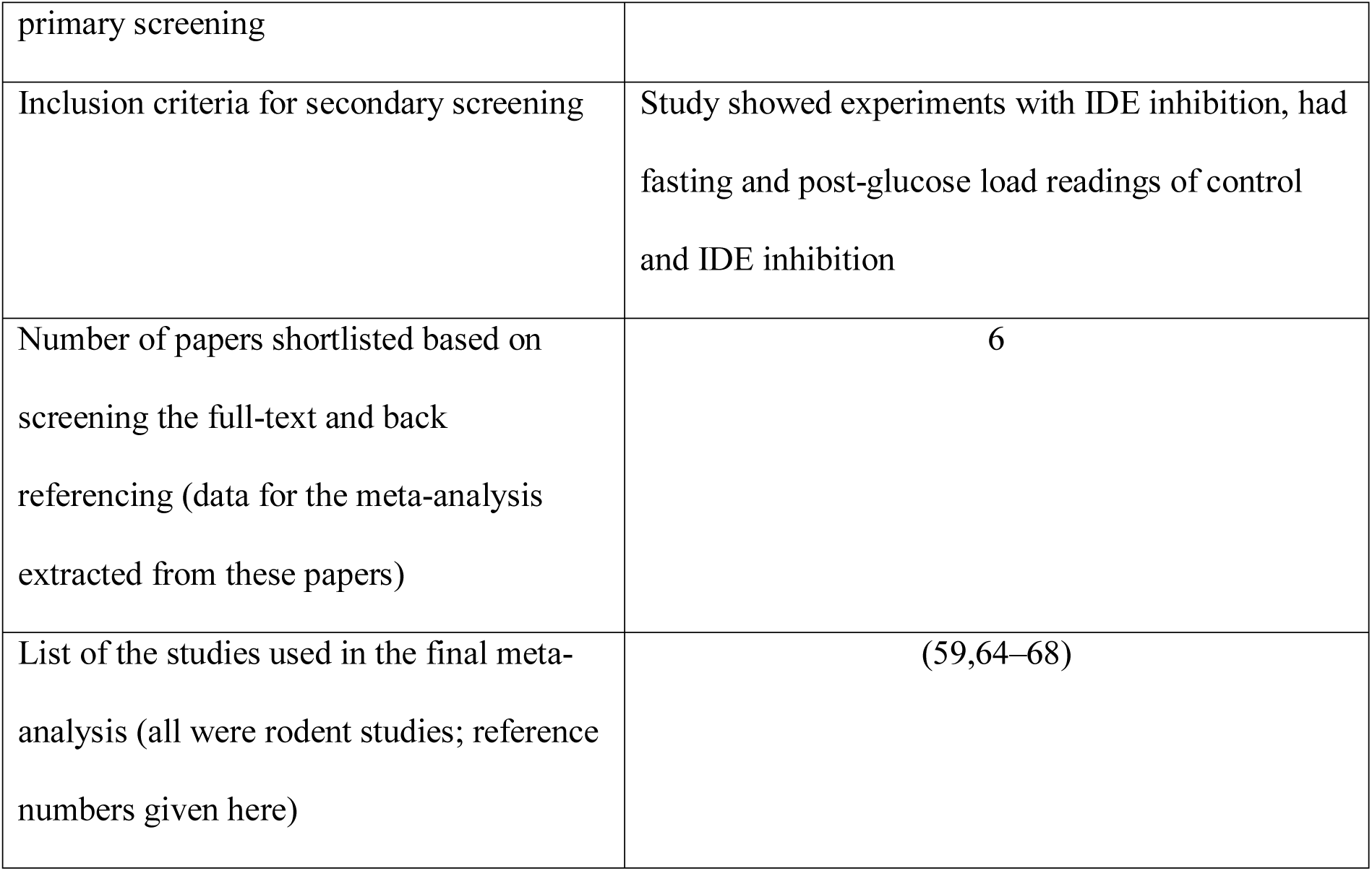
Systematic literature review for studies on insulin degrading enzyme inhibition/knockout (meta-analysis 2).

#### 3.1.2 Decrease in insulin

We performed a systematic literature review for experiments in which there was sustained suppression of insulin production. Two insulin suppressing agents have been repeatedly used to lower insulin production in rodent models as well as in humans.

i. Diazoxide (DZX): Diazoxide is a potassium channel activator which causes reduction in insulin secretion by the β hyperpolarized state by opening the channel (61). It has been used as a drug to modulate insulin secretion for research and therapeutic purposes (62).
ii. Octreotide (OCT): Octreotide is a somatostatin analogue which inhibits insulin and growth hormone. It has been used to reduce insulin secretion *in vitro* and *in vivo* (63).

We searched the literature systematically for studies where the insulin levels have been altered using either DZX or OCT and glucose tolerance has been examined using a GTT after DZX/OCT treatment (table 4). It should be noted that this literature includes a significant proportion of human trials. We also searched literature for studies in which insulin was suppressed by other methods.

**Table 4:**
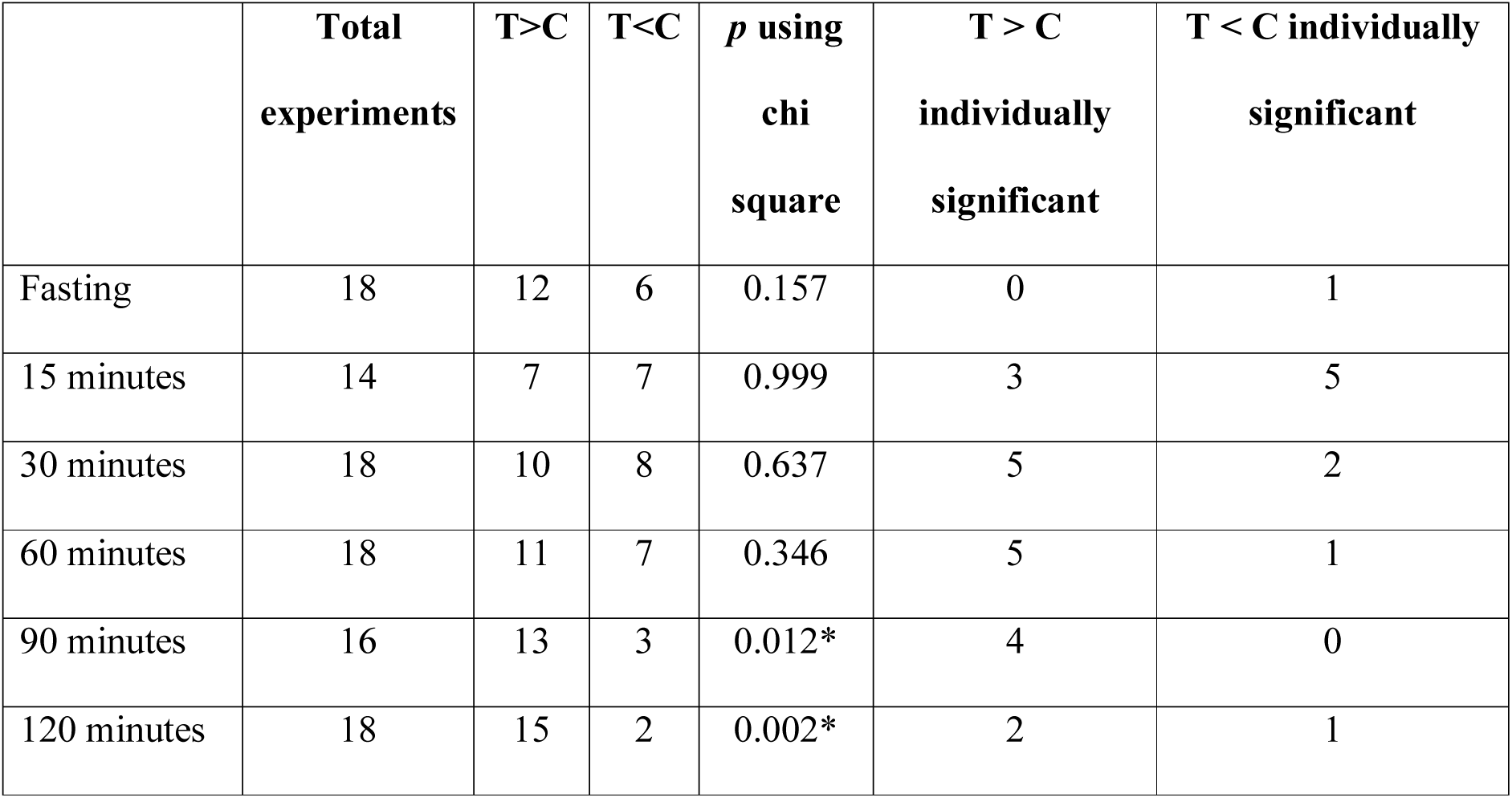
Comparison between steady-state (fasting) and perturbed-state (post glucose load) of control and IDE suppression.

**Table 5:**
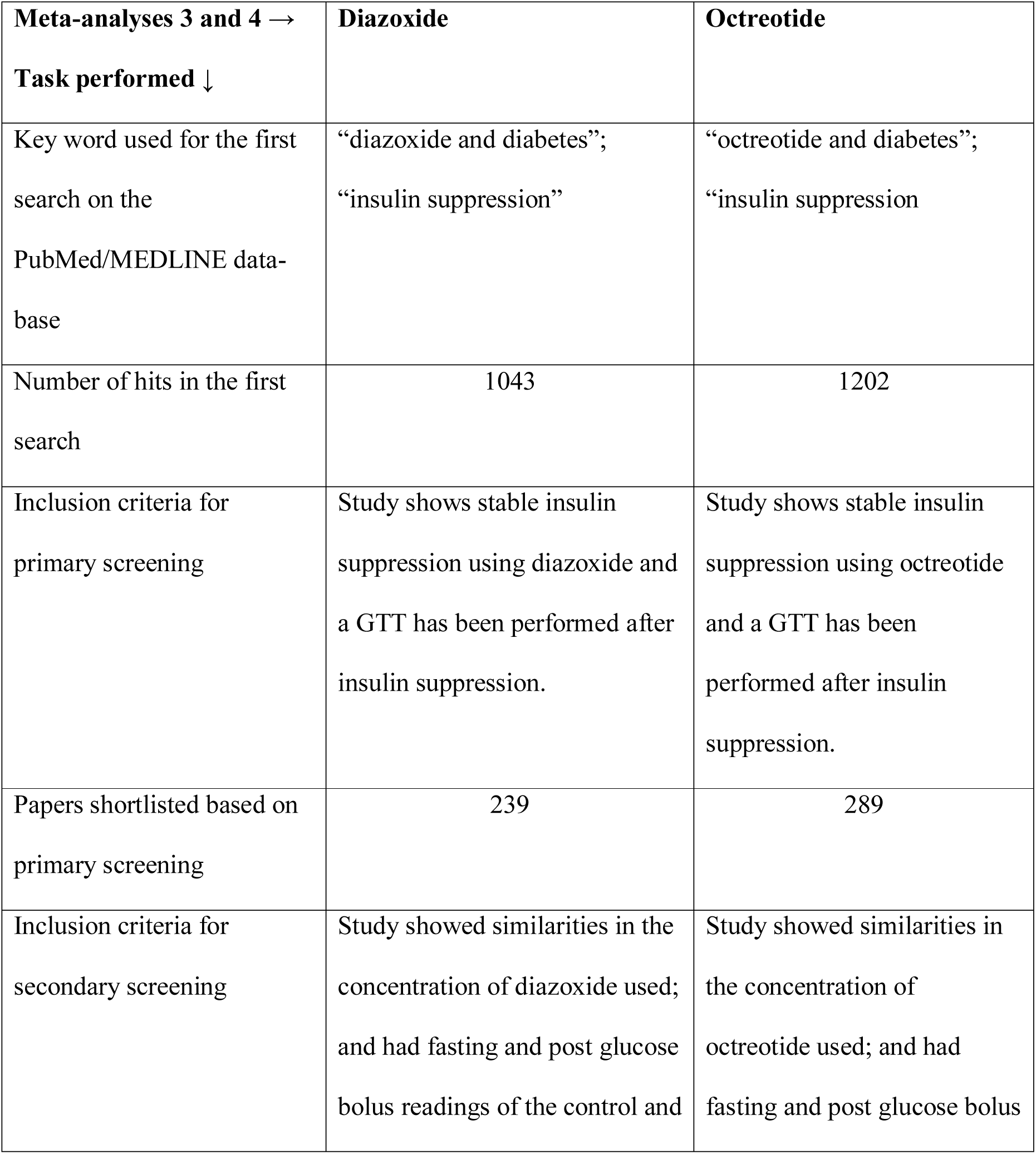
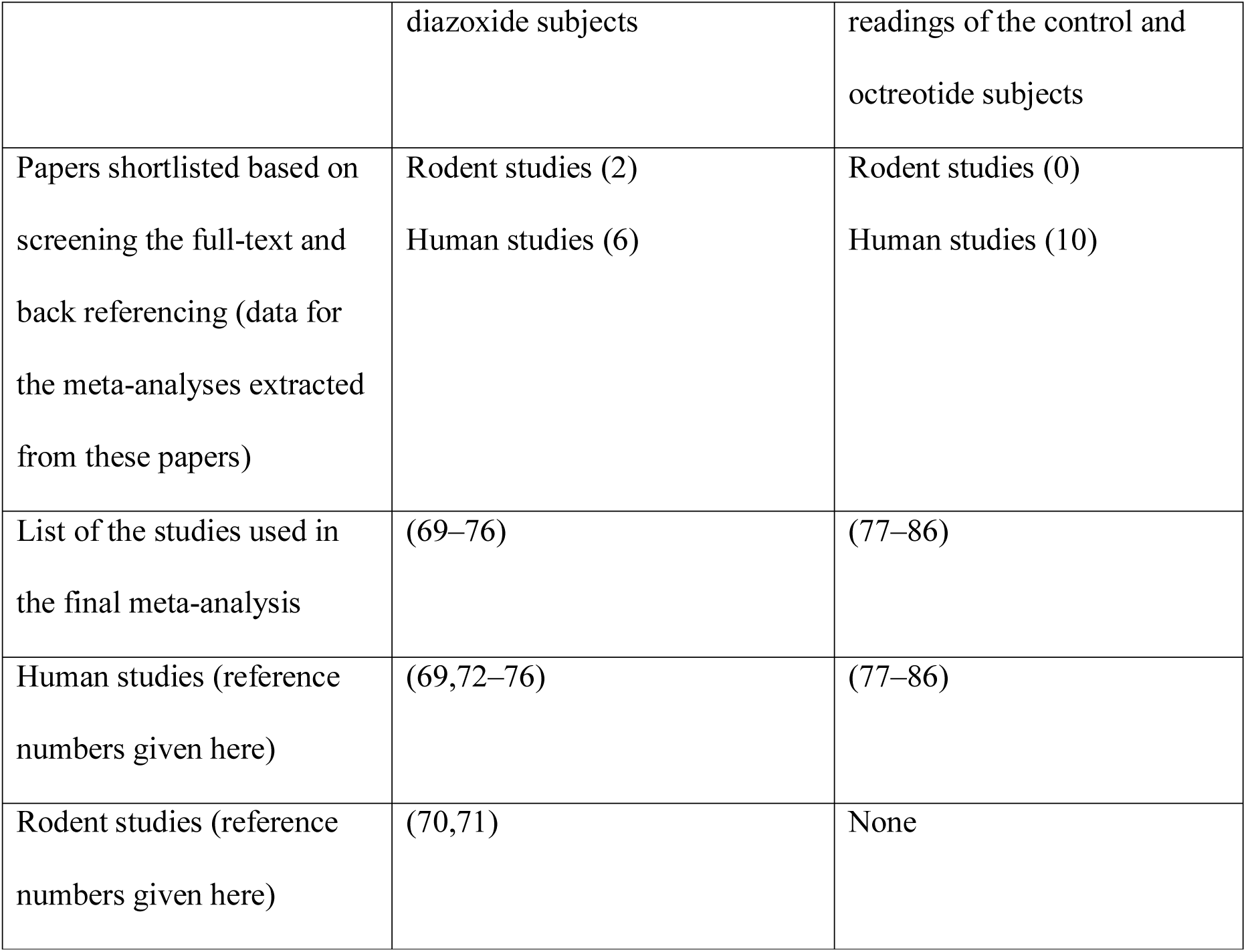
Systematic literature review for studies on insulin suppression with diazoxide and octreotide

### 3.2 Results

#### 3.2.1 Increase in insulin by suppression of IDE

We found 6 publications that described 18 experiments that allowed comparison of GTT between raised insulin groups and control group (table 2 of supplementary information 1). Meta-analysis revealed no significant difference in the fasting glucose. In only one out of 18 experiments the treatment group had lower fasting glucose than the control. During the GTT curve, at 90 and 120 minutes the difference between treatment and control were significant but in the opposite direction of the expectation. While rise in insulin level should reduce plasma glucose, it increased in 15 out of 18 studies, two of which were individually significant and the difference was significant in non-parametric meta-analysis. Across all time points along the GTT, the plasma glucose in the treated group was greater than the control group in majority of the experiments. Thus, in this class of experiments increasing insulin failed to reduce glucose at the steady-state as well as post glucose load.

#### 3.2.2 Decrease in insulin: Suppression by diazoxide or octreotide

We found 8 papers describing 14 experiments for diazoxide treatment and 10 papers with 15 experiments for octreotide treatment (tables 3 and 4 from supplementary information 1 respectively). It can be seen from table 6 and fig 6 that for both of the insulin suppressing agents, suppression of insulin did not result into increased fasting glucose. Further at 120 minutes post glucose load there was a marginally significant rise in glucose in the insulin suppressed group as compared to control group. This demonstrates that pharmacological suppression of insulin was unable to raise plasma glucose level in a fasting steady state. There was inconsistent but significant rise post glucose load.

**Table 6:** Comparison between steady-state(fasting) and perturbed-state (post glucose load) of control and insulin suppression.

|  | **Total**<br>**studies** | **T>C** | **T<C** | ***p* using**<br>**chi**<br>**square** | **T > C individually**<br>**significant** | **T < C individually**<br>**significant** |
| --- | --- | --- | --- | --- | --- | --- |
| **Diazoxide treatment** |  |  |  |  |  |  |
| Fasting | 14 | 10 | 3 | 0.052 | 5 | 0 |
| 15 minutes | 7 | 6 | 1 | 0.059 | 1 | 1 |
| 30 minutes | 12 | 10 | 2 | 0.021* | 2 | 1 |
| 60 minutes | 13 | 9 | 4 | 0.166 | 4 | 2 |
| 90 minutes | 3 | 3 | 0 | 0.083 | 2 | 0 |
| 120 minutes | 14 | 10 | 3 | 0.052 | 6 | 1 |
| **Octreotide treatment** |  |  |  |  |  |  |
| Fasting | 15 | 6 | 7 | 0.781 | 0 | 0 |
| 30 minutes | 14 | 4 | 10 | 0.108 | 0 | 2 |
| 60 minutes | 14 | 4 | 10 | 0.108 | 2 | 1 |
| 90 minutes | 13 | 5 | 8 | 0.405 | 1 | 0 |
| 120 minutes | 15 | 7 | 8 | 0.797 | 1 | 1 |

**Figure 1:**
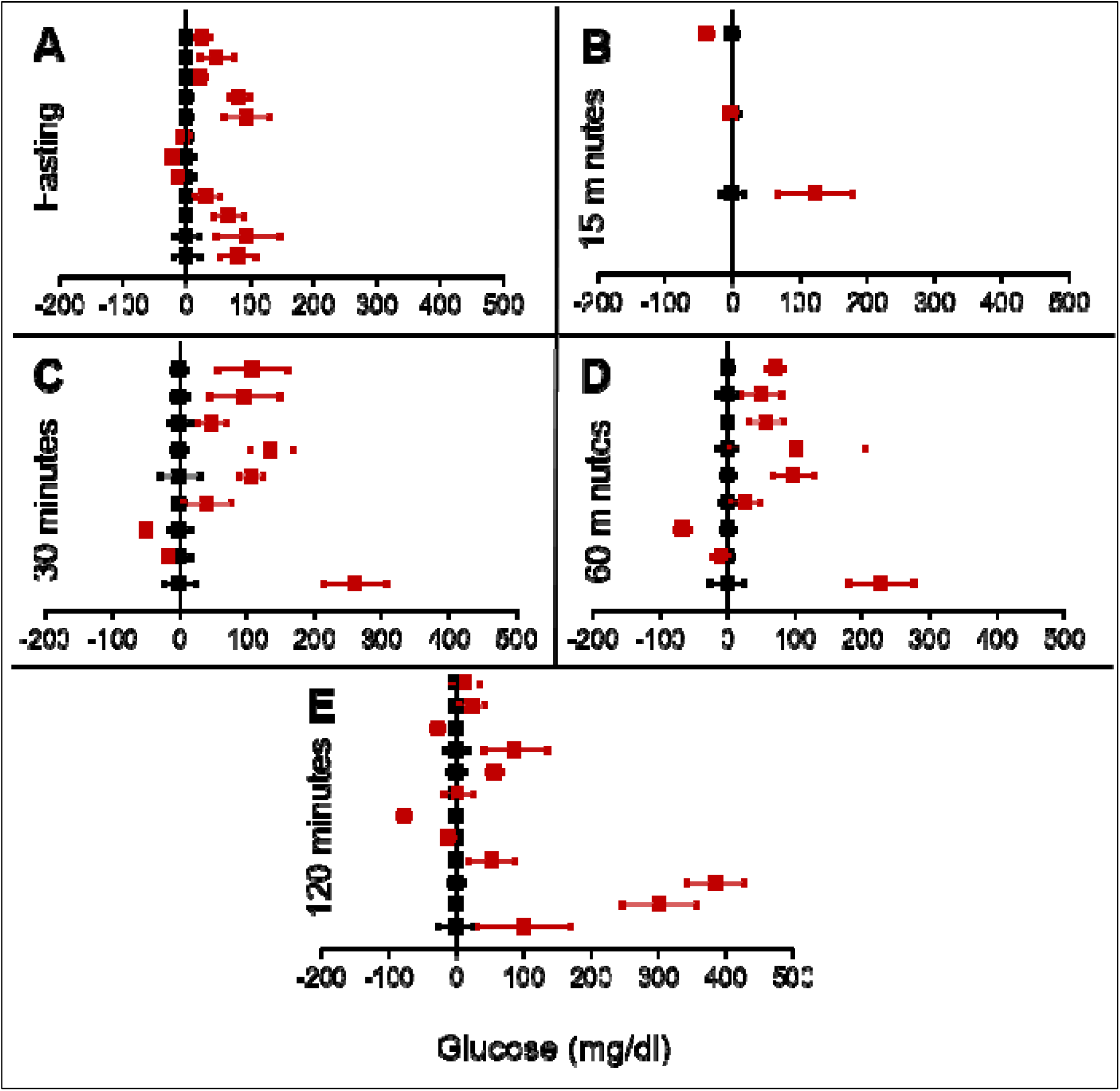
Glucose levels for control (black squares) and FIRKO (red squares) at steady-state and perturbed state. The forest plot is normalized to the control and difference of FIRKO glucose levels plotted with ± 95% CI. fasting glucose and (B) to (E) post glucose load glucose at different time intervals.

**Figure 2:**
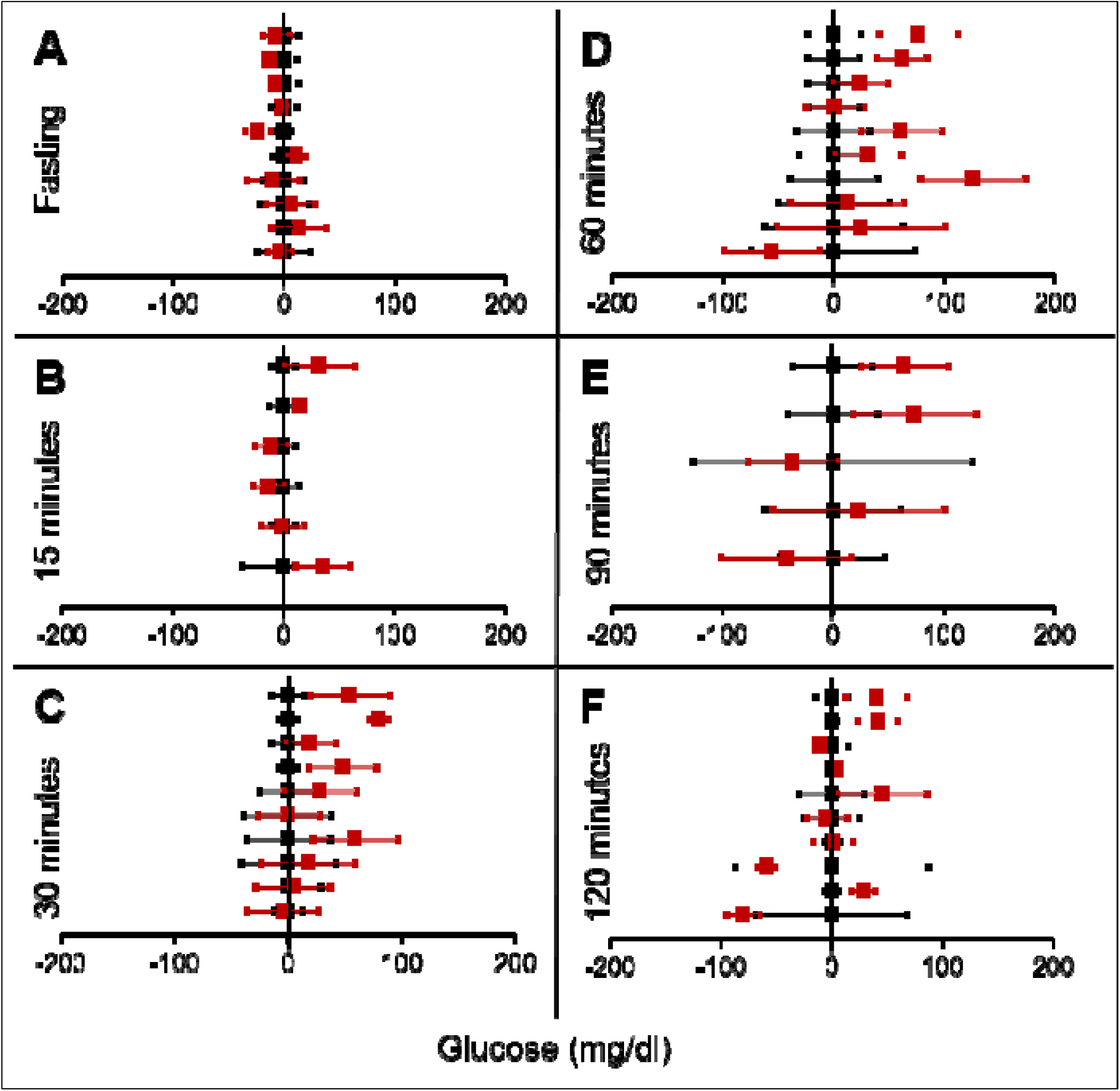
Results with **MIRKO** represented as in figure 1. Note that fasting glucose does not differ from the control in any of the experiments.

**Figure 3:**
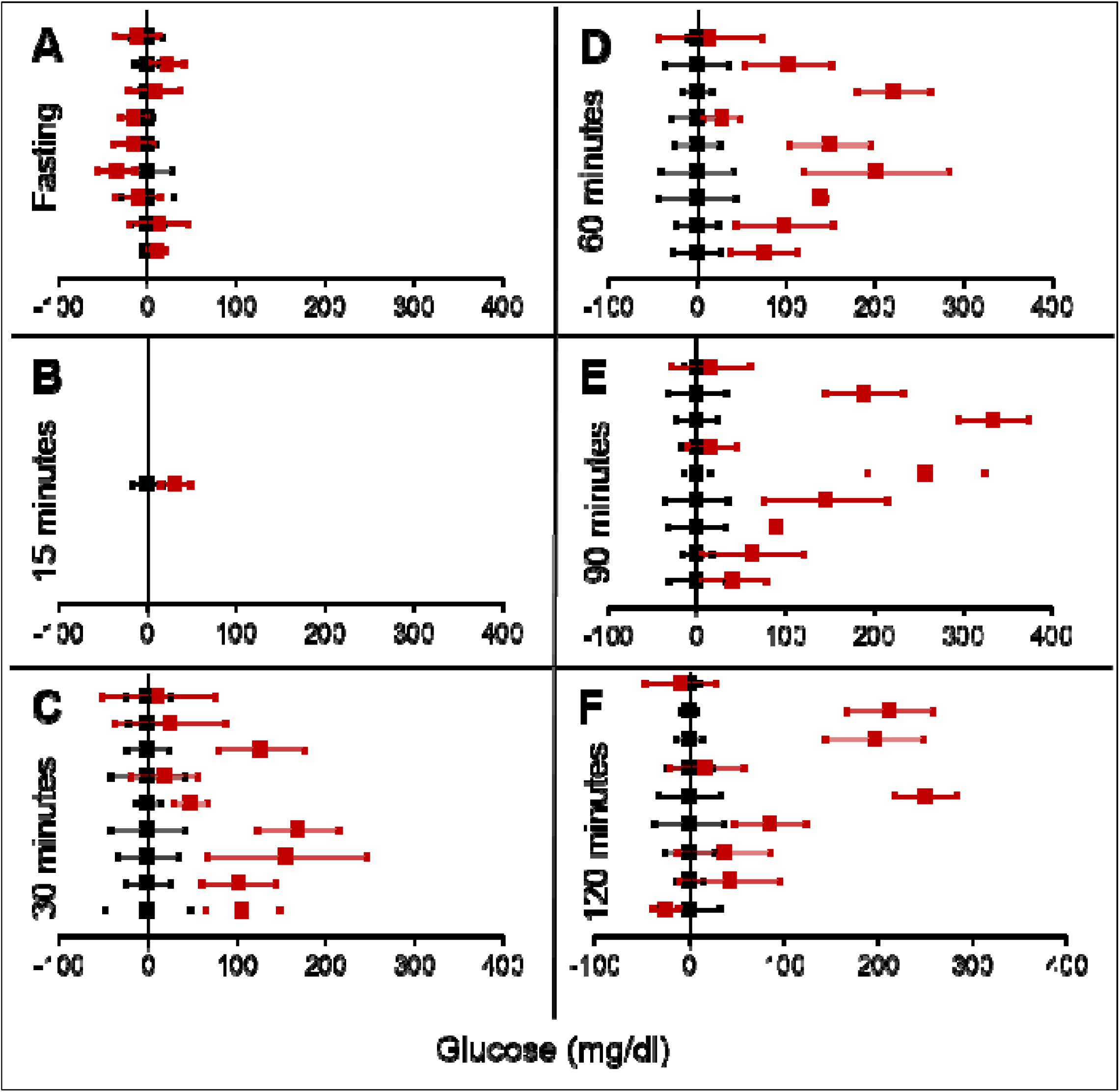
Results with **LIRKO** represented as in figure 1. Note that inconsistent with classical belief, liver specific insulin receptor knockout does not show significant effect on fasting glucose in any of the experiments. On the other hand post load glucose is consistently higher.

**Figure 4:**
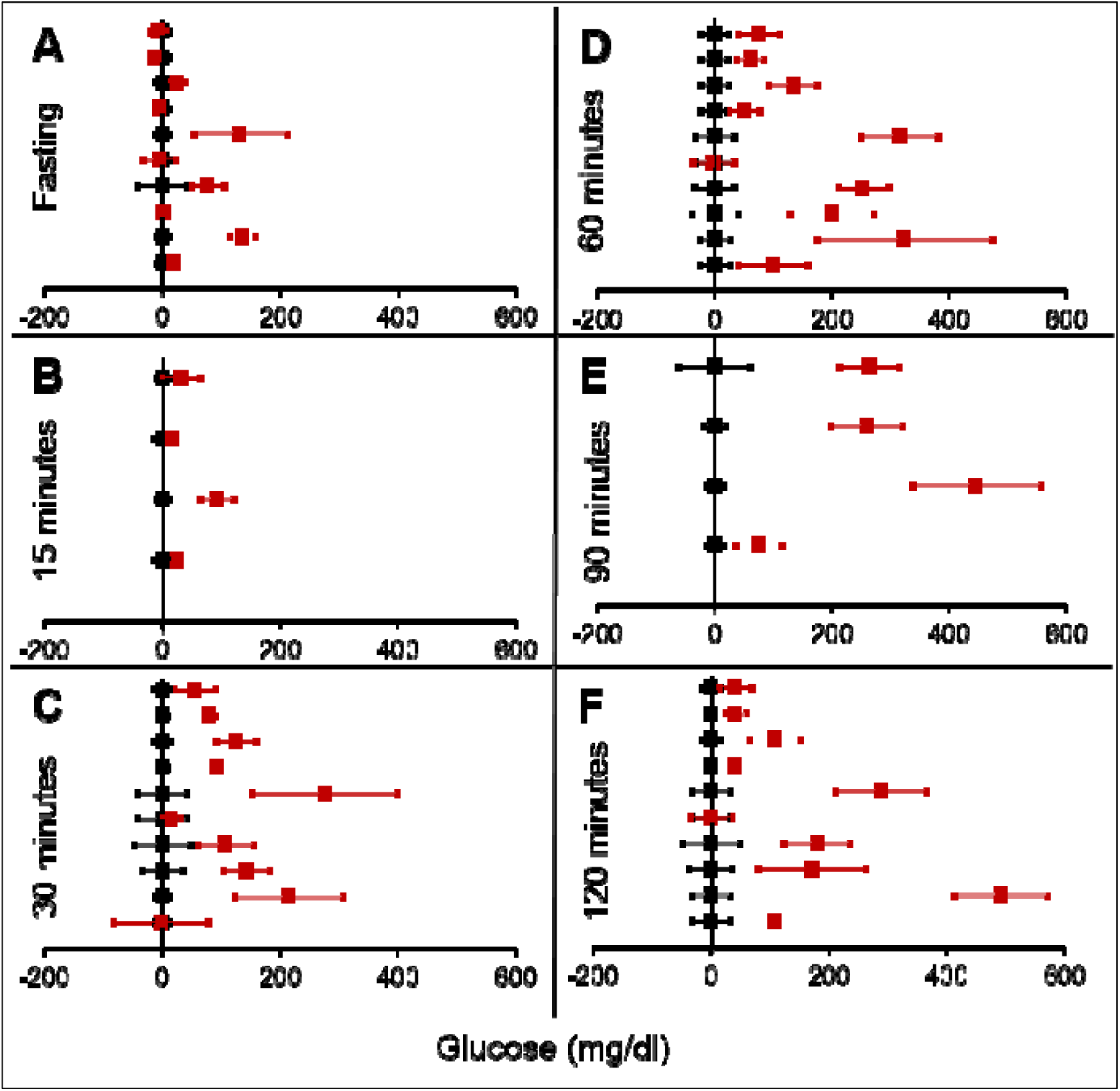
Results with **βIRKO** represented as in figure 1. Note that fasting glucose does not differ from the control in any of the experiments.

**Figure 5:**
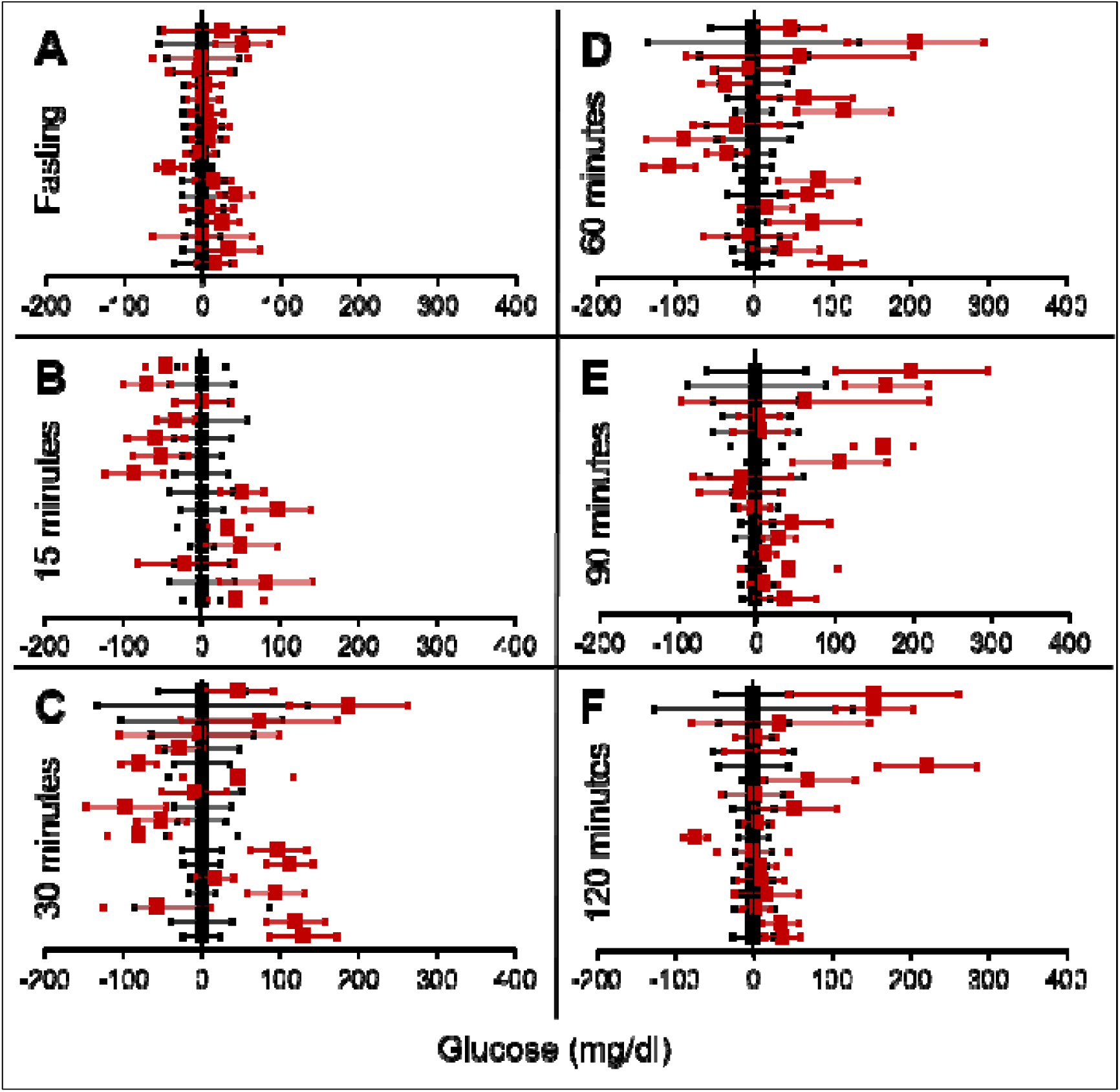
Results with **IDE inhibition** represented as in figure 1. Note that fasting glucose does not differ significantly from the control. At 90 and 120 minutes the trend is higher mean glucose than control which is contrary to the expectation in an experiment with sustainable rise in insulin.

**Figure 6:**
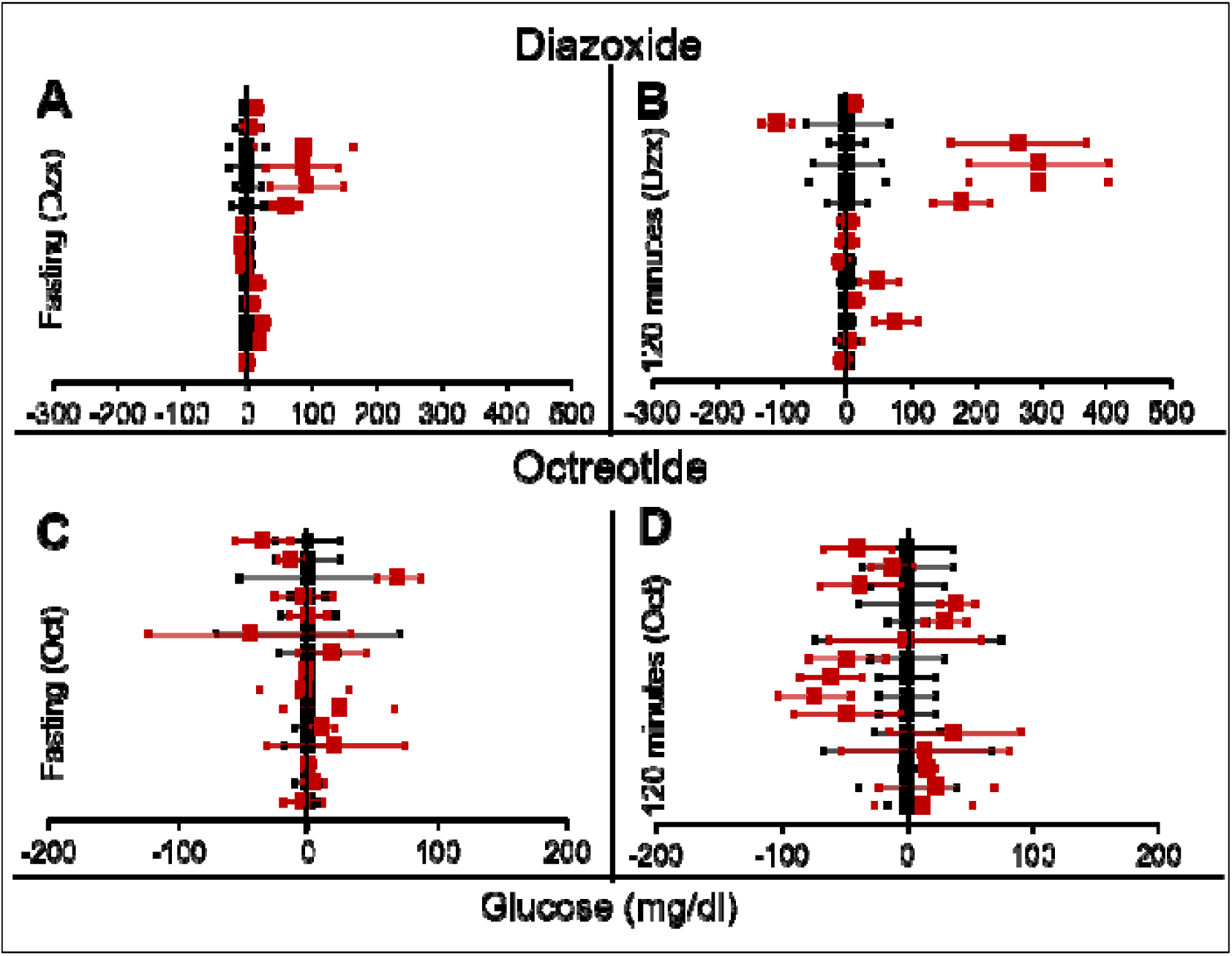
Results with **Insulin suppression with Diazoxide and Octreotide** represented as in figure 1. Note the inconsistencies across studies.

We found more means of insulin suppression in which GTT after suppression was reported, but there were not many published replications of the experiments coming independently from different research groups. Therefore meta-analysis was not warranted. We briefly review their results here.

*Suppression by Protein restriction:* Dietary protein deprivation is another method of insulin suppression. This also led to a decrease in plasma insulin levels; however fasting glucose levels did not increase (87).

*Suppression by insulin siRNA*: Transgenic mice for insulin-siRNA along with IDE overexpression, showed decreased levels of insulin. Again the fasting glucose levels remained normal while there was a change in glucose tolerance curve (figure 7) (88). The curves in figure 7 are typical of insulin receptor knockout or insulin suppression experiments where, in individuals with impaired insulin signalling the glucose peak is higher which returns to steady-state much later than the controls, but the fasting steady-state level is not different.

**Figure 7:**
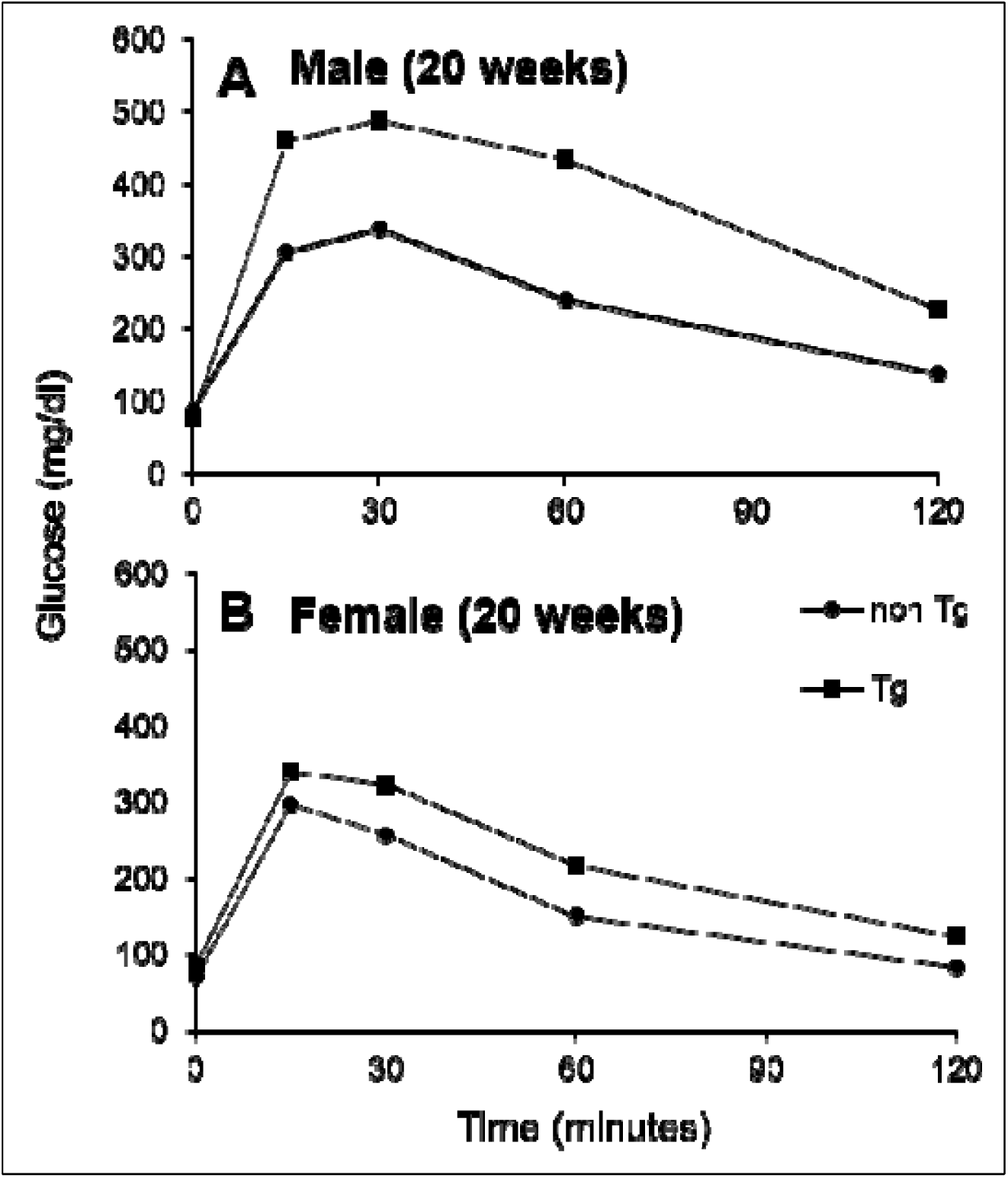
Intra peritoneal glucose tolerance test (panel A and B). Fasting glucose levels in both the siRNA treated and untreated group remain unaltered in male and female mice. 15 minutes after the glucose injection, the treated mice show higher glucose levels relative to the untreated mice and this effect is seen throughout till 120 minutes. Figure redrawn from data by (88).

*Suppression of insulin by partial gene ablation:* In rodents, there are two insulin genes *Ins1* and *Ins2* (89). A double knockout of both the genes results in death, but ablation of either of the genes does not alter the glucose tolerance significantly suggesting redundancy (90). There are studies in which one gene is completely knocked out and the other one is a heterozygote (90–94). Reduced insulin gene dosage did not consistently result into fasting hyperglycemia in these studies although it offered protection against some of the effects of hyperinsulinemia.

## 4. Steady-state versus post feeding glucose in STZ rats

Streptozotocin (STZ) induced diabetes is a popular model of rodent diabetes. STZ acts by specifically destroying the insulin producing β cells of the pancreatic islets. A low dose of STZ that destroys a substantial population of β cells but does not lead to total destruction of their population is often perceived as a good model for T2D, whereas a high dose of STZ that destroys the β cell population almost entirely is perceived as a model of T1D. We searched literature to look for studies that carefully differentiated between steady-state glucose from post load glucose in STZ models but did not find any studies that make this distinction clear. Therefore, we designed and conducted experiments to differentially study the steady-state and perturbed-state glucose levels in rats treated with STZ.

### 4.1 Experimental methods

#### 4.1.1 Animal model and conditions

The experiments performed on Sprague Dawley (SD) rats had been approved by the Institutional Animal Ethics Committee at IISER, Pune (Protocol Number IISER/IAEC/2016-02/006) constituted by CPCSEA, Govt. of India. All the rats were housed in a facility with a temperature of 23±2°C and a 12-hour light/dark cycle with standard rat chow and water available ad libitum. The bedding of the cages was changed every three days.

#### 4.1.2 STZ treatment for insulin suppression

Male, SD rats weighing 180-200 g were injected with STZ at 50 mg/kg body weight. The STZ was dissolved in Citrate Buffer (Citric Acid: 0.1M and Sodium Citrate: 0.1M). Injection of citrate buffer alone was used as control.

#### 4.1.3 Fasting and post-feeding glucose in 12 day follow up

Three days after the STZ injection, the rats were fasted for 16 hours and glucose was measured using the hand held Accu-Chek Glucometer. The rats were then given 40 grams of Standard Chow for 8 hours. Food was weighed and post-meal glucose was measured after three hours. The protocol was repeated for 12 days and body weight, food weight and glucose readings were taken daily. 12 animals per group were used for this experiment.

#### 4.1.4 Duration of fasting

An experiment was also performed to see how much time was required to reach a steady-state of glucose after removal of food. The food was removed from the STZ and Control animals after ad libitum availability and glucose readings were taken after 3 hours, 6 hours, 9 hours, 12 hours and 16 hours. After a recovery of three days, glucose levels were measured only at 16 hours after removing the food. 9 STZ treated animals and 10 Control animals (injected with citrate buffer) were used for this experiment.

### 4.2 Results

Among the STZ treated rats, all the animals showed significantly higher post load glucose than the control group on all the 12 days sampled. However, in 10 out of the 12 days the 16-hour fasting glucose was not significantly different from the control although the variance was substantially greater than that of the control (figure 8).

**Figure 8:**
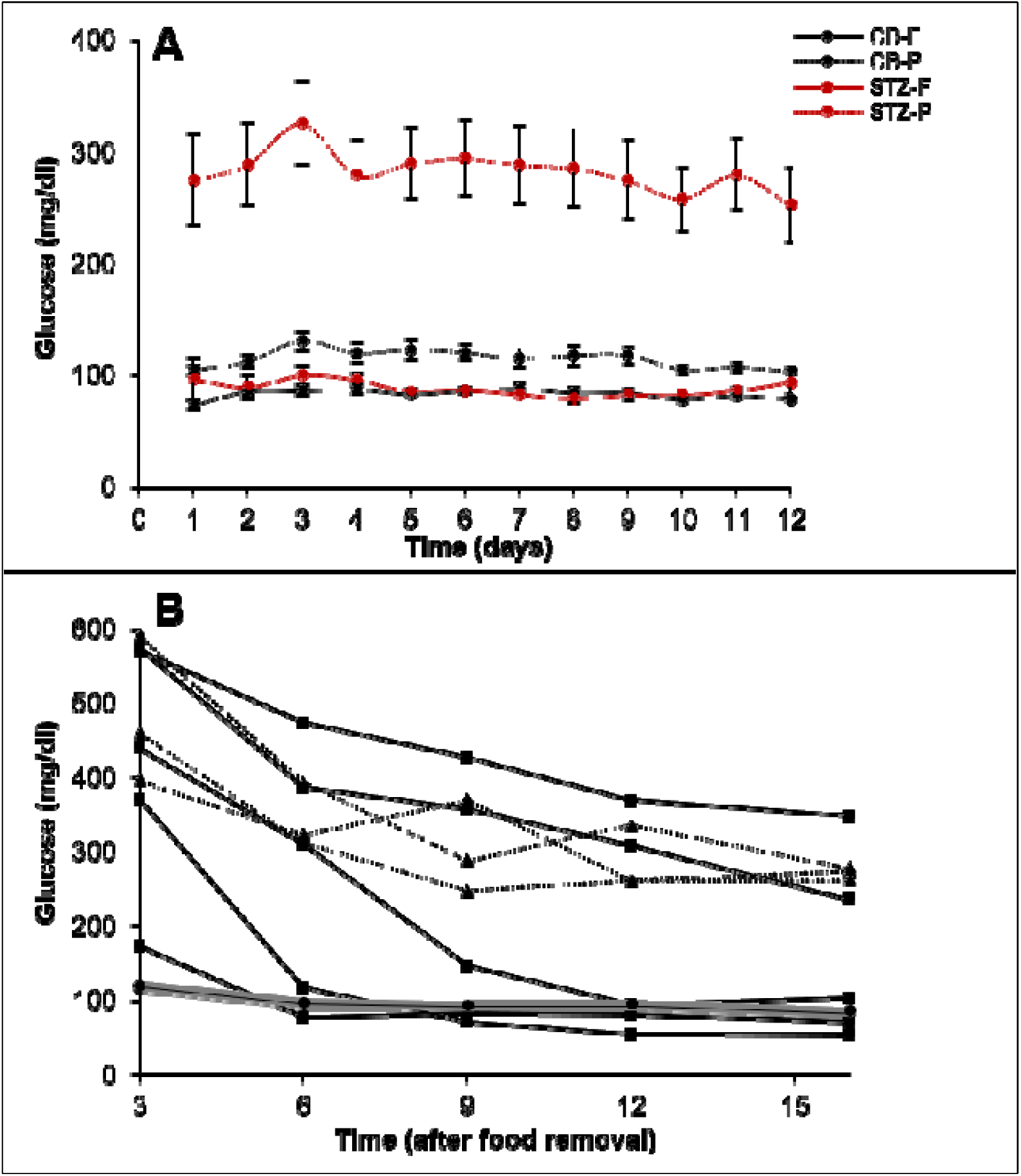
Treatment of SD rats with STZ. (A): The 16-hour fasting and post-meal glucose values of treated (STZ 50mg/kg) and control rats (Citrate buffer CB) over 12 days. N= 12 for each group. Note that on 10 out of 12 days the mean fasting glucose of the treated group (STZ-F) was not significantly different from the control mean (CB-F). Post feeding the treated group (STZ-P) has substantially greater mean than the control (CB-P). (B)Time course of glucose during 16 hours fasting. X axis represents the time after removal of food when the glucose readings are taken and Y axis represent the glucose levels. The grey band represents the upper and lower bounds of 95% CI of the control group with the mean glucose values represented by filled circles. Filled squares represent individuals that showed a monotonic decrease in glucose levels. In three animals the glucose levels reduced at or below the control levels and in two others they showed a continued monotonic decrease but did not reach the normal level in 16 hours. Filled triangles with dotted lines represent the individual time courses of the three STZ treated rats which showed some indications of stabilizing at a steady-state above the normal.

A close look at the time course of fasting in the two groups revealed that in 4 out of 9 STZ animals the glucose levels reached the normal range but with substantial delay as compared to control animals. In two more animals the levels did not reach the normal range till 16 hours but a monotonic decrease continued throughout the period, indicating that their blood glucose may not have reached a steady-state in 16 hours. Only in 3 animals the 16-hour glucose was higher than the control range with some indications of stabilizing at a higher level. In the time course experiment, the animals are handled frequently leading to some unavoidable stress. In the 12 day follow up plasma glucose is estimated only after 16 hours and here there is no significant difference in the control and STZ animals on 10 out of 12 days. Furthermore the individuals that showed higher 16 hour fasting glucose did not do so consistently. In the 12 days follow up, the distribution of 16-hour fasting glucose was typically skewed with one or two outliers having high glucose levels. Interestingly the outliers were not the same animals every day. There was considerable day to day variation in individuals and averaged over the 12 days, none of the STZ animals showed significantly higher fasting glucose than the controls although they consistently showed higher post feeding glucose.

Thus, these experiments show on the one hand that STZ treatment failed to increase steady state glucose levels significantly and consistently. On the other, the STZ animals took substantially longer and rather unpredictable time to reach a steady-state and even at 16-hours of fasting, all individuals need not have attained a steady state. These results warrant caution against considering fixed hours fasting glucose as steady-state glucose in experimental or epidemiological data. While in healthy individuals it is well established that following glucose load a steady-state is regained in about two hours, it is possible that in experimental impairment of insulin signalling or in clinical diabetes, plasma glucose takes substantially longer time to reach a steady-state and overnight fasting need not represent a steady-state in all cases.

## 5. Theoretical and mathematical considerations

In this approach we elaborate on the theoretical underpinnings of insulin-glucose relationship. We also explore possible explanations for the unexpectedly consistent failure of experimental insulin signal impairment to alter steady-state glucose level. Simultaneously we make differential predictions from alternative homeostasis models that can be tested in human epidemiological data.

### 5.1 Choice of Models for glucose homeostasis

The fasting state has been generally accepted to be a steady-state for glucose concentration for several reasons. In a given healthy individual the fasting glucose levels are stable in time (95, 96). The post-meal peak of glucose and insulin returns to the fasting level within a few hours and remains stable over a long time. The fasting state is considered and modelled as a steady-state by the widely used HOMA model (28, 29). Classically the negative feedback loops are assumed to work through insulin and insulin is taken as a determinant of steady-state glucose level. Most popular models of glucose homeostasis work on this assumption although non-steady-state models of insulin resistance exist (97).

A critical question in glucose homeostasis is whether the fasting steady-state glucose level is a consequential balance between glucose production and glucose utilization rates (consequential steady-state CSS) or whether there is a target glucose level that is maintained by sensing and correcting any changes in it (targeted steady-state TSS). The difference in the two can be visualized by the tank water level analogy (fig 9). If a tank has an input tap releasing water in it at a constant rate and has an outlet at the bottom through which water escapes proportionate to the pressure of the water column, a steady-state is invariably reached. The steady-state level is decided by the rate of intake and the size of the outlet. This is a CSS which will change with any change in the size/capacity of the input or the outlet tap. In contrast to CSS, in a TSS there is a desired water level and sensors are placed above and below the desired level such that when the level goes below the lower sensor the input is switched on or its rate increased and/or output switched off or its rate decreased.

**Figure 9:**
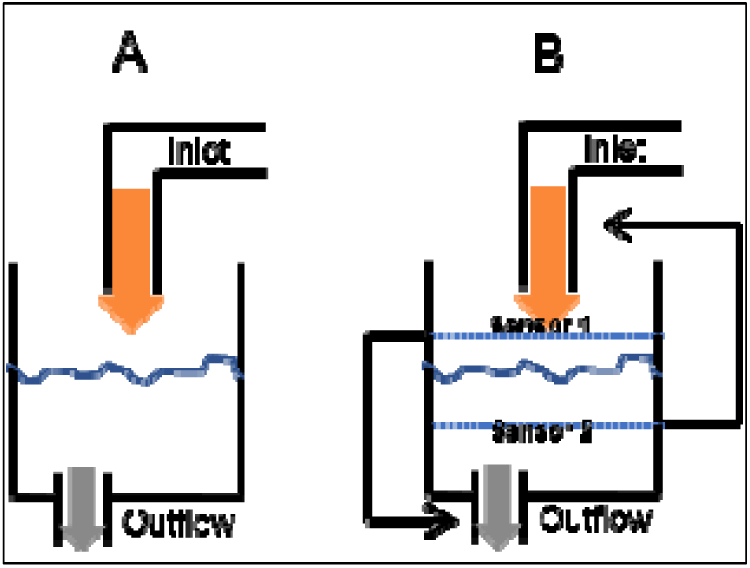
The consequential steady-state (CSS) (A) and targeted steady-state (TSS) (B) models of homeostasis illustrated with a tank water level analogy. In CSS a change in the size of inlet or outlet tap, analogous to insulin sensitivity can change the steady-state level. In a TSS model, a change in the tap size will alter the time required to reach a steady-state but will not change the steady-state level.

In glucose homeostasis, in a fasting state, liver glucose production is analogous to the inlet tap and tissue glucose uptake analogous to the size of the outlet, both being a function of insulin signalling. Most models of glucose regulation assume CSS (28,29,58,98–100) (97). It has not been critically examined whether CSS or TSS describes glucose homeostasis more appropriately. This is important because if TSS model is appropriate, insulin resistance and relative insulin deficiency will not result into altered steady-state glucose levels although the time required for reaching a steady-state after perturbation might change. If CSS model is appropriate, insulin resistance or altered insulin levels are bound to change fasting glucose levels. The failure of insulin receptor knockouts and insulin suppression experiments to alter the fasting steady state, along with the delay in reaching the steady-state indicates that TSS model is likely to describe glucose homeostasis more appropriately. The TSS model requires mechanisms of sensing any departure from the targeted steady state. Such mechanisms are not known in peripheral systems but glucose sensing neurons are certainly known to be present in the brain. Therefore, if TSS is a more appropriate model, the CNS mechanisms are likely to be central to glucose homeostasis, particularly in determining the steady-state levels; whereas insulin signalling would play a role in determining the rate at which a steady-state is reached after perturbation.

It is possible to make other testable predictions of TSS and CSS models. In the normal healthy individual, increased glucose utilization is expected to decrease fasting glucose levels by the CSS model but not by the TSS model. Human experiments have shown that sustained exercise does not reduce plasma glucose, in fact it might increase (101). In order to match with experimental data, CSS based models of glucose dynamics during exercise need to include additional terms which involve neuronal mechanisms such as direct stimulation of liver glucose production in response to exercise through sympathetic route (102). This brings the model close to a TSS model. If TSS model describes glucose homeostasis more appropriately, reduced insulin signalling is not expected to change steady-state glucose but only alter the time course to reach a steady state.

The mechanism of attaining a hyperinsulinemic normoglycemic prediabetic state is different by the CSS and TSS models. By the classical CSS based pathway, obesity induced insulin resistance is believed to be primary. The insulin resistance reduces glucose uptake and the excess glucose triggers a compensatory insulin response. The resultant hyperinsulinemia compensates for insulin resistance keeping the fasting glucose levels normal. Detailed analysis of the model and matching its prediction with empirical data has refuted this model (30). One of the intuitively appealing reasons for this refutation is that after the heightened insulin levels normalize glucose, there is no reason why insulin levels remain high.

Therefore, a steady-state with hyperinsulinemia and normoglycemia is impossible by the CSS model but it exists in a prediabetic state. If a “compensatory” insulin response is mediated by glucose, one would expect a positive correlation between fasting glucose (FG) and fasting insulin (FI) and no correlation between insulin resistance and β cell responsiveness.

By the TSS model, on the other hand, compensatory response is possible in either way. Primary insulin resistance may increase the glucose levels transiently, but when glucose sensing mechanisms detect the change a compensatory response can be operational. By this mechanism a hyperinsulinemic normoglycemic state is possible. Alternatively, primary hyperinsulinemia (7,103–105) can also be compensated by increased insulin resistance by hitting the lower level of sensing which would trigger compensatory insulin resistance. Even in this case a hyperinsulinemic normoglycemic state is possible. Both glucose sensing neurons and neuronal regulation of insulin release and liver glucose production are well known. In the compensatory response mediated by TSS pathways there need not be a correlation between fasting insulin and fasting glucose, but insulin resistance and β cell response would be correlated.

Also using a simple CSS model (see supplementary information 2 for details), simulations show that the correlation coefficient and regression slope in the insulin-glucose relationship would remain the same in the fasting as well as post-meal state although the range of glucose and insulin levels will be different. On the other hand, in a TSS model the post-meal glucose and insulin levels are expected to be correlated but the steady-state levels may not. We test these predictions by the alternative models using human epidemiological data below.

We argued above that since on impairment of insulin signalling, the time required to reach a steady-state can be substantially longer, overnight fasting may not ensure a steady-state in all individuals. Fasting hyperglycaemia in T2D can have two alternative (but not mutually exclusive) causes. Either it represents the failure to reach a steady-state in the specified fasting period, or it is because of mechanisms other than reduced insulin action. The TSS model can make differential predictions from the two alternative causes since it predicts a positive correlation between plasma glucose and plasma insulin in the post-meal state but loss of this correlation on reaching a steady state. In population data, if some individuals have reached a steady-state but a few others haven’t we would expect a correlation significantly weaker than the post-meal correlation. These predictions can be tested in epidemiological data.

## 6. Analysis of insulin glucose relationship in steady and perturbed-state in human data: Epidemiological inquiry

Here we use human epidemiological data to test the correlational predictions made by the CSS versus TSS models of glucose homeostasis.

### 6.1 Methods

#### 6.1.1 Epidemiological data

The three data sets used here come from two different studies: (i) Coronary Risk of Insulin Sensitivity in Indian Subjects (CRISIS) study, Pune, India (106) and (ii) Newcastle Heart Project (NHP), UK (107). Data from the latter is divided into two groups as the subjects belong to different ethnicities namely European white and south Asian and we will prefer to analyse the two groups separately since certain ethnic differences are likely to be present in the tendency to develop metabolic syndrome (108, 109). Hence all the comparison of predictions with the data has been done independently for the three data sets. All the studies are population surveys that include non-diabetic (fasting glucose values less than 110mg/dl) and diabetic individuals (fasting glucose values above 110 mg/dl) and the clinical history, morphometric parameters, glucose and insulin during fasting and oral glucose tolerance test (OGTT) of the subjects were recorded. In the analysis below we included only the non-diabetic groups in which the homeostatic mechanism can be assumed to be intact and therefore any hypothesis about it can be tested. Most of the individuals in the diabetic group would be under different drug regime affecting glucose-insulin dynamics in different ways and therefore we exclude that group for the analysis.

#### 6.1.2 Analysis

Linear regression and correlation were used to compare the glucose-insulin relationship in steady-state (fasting) versus perturbed-state (post glucose load) in the three data sets along with the relationships between HOMA-IR and HOMA-β derived from the fasting data.

### 6.2 Results

In all the three data sets there was weak (R^2^ range 0.017 to 0.057) but significant correlation between fasting glucose (FG) and fasting insulin (FI) and strong correlation between HOMA-IR and HOMA-(R^2^ range 0.20 to 0.83) (figure 10).

**Figure 10:**
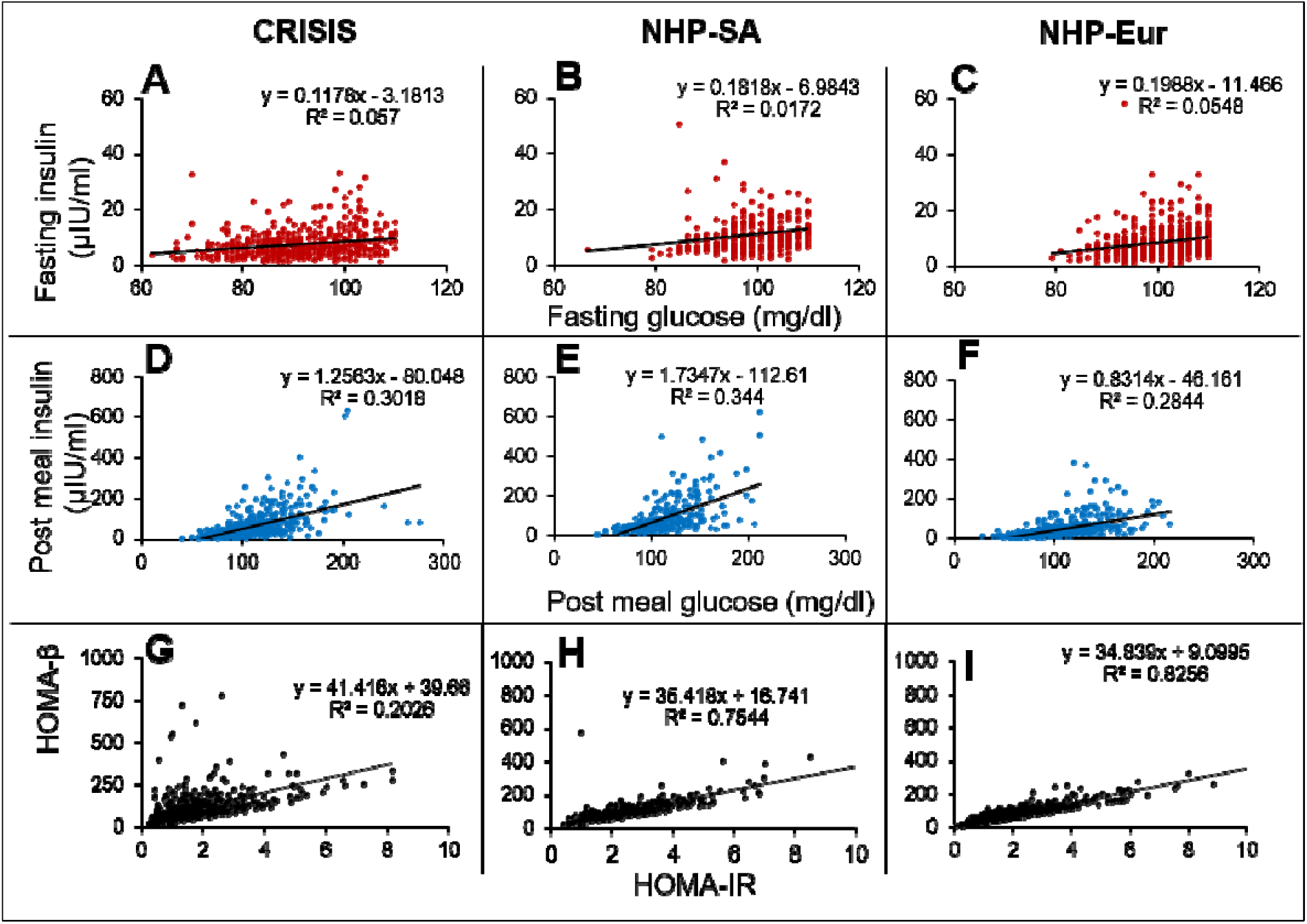
The Fasting Glucose-Fasting Insulin, Post-meal Glucose-post-meal Insulin and HOMA-IR HOMA-β scatter plots in non-diabetic populations in the three data sets. The FG-FI correlation is weak as compared to post-meal correlation. The HOMA-IR and HOMA-β correlations are very strong in all the three data sets, which is not expected by the classical insulin resistance theory.

It was seen in all three data sets that the correlation coefficients for glucose and insulin were an order of magnitude higher in the post-meal cross sectional data than in the fasting state (Table 7 and fig 10). Also, the regression slopes in the post-meal data were substantially different from fasting data unlike what is expected by the CSS model (Supplementary information 2). By both the sets of predictions the CSS model predictions are rejected. The HOMA-IR HOMA-β correlation, as well as the difference between the regression correlation parameters between fasting and post-meal data are compatible with predictions of the TSS model. However, although weak, there is significant correlation between FG and FI unlike what may be expected by a steady-state TSS model. This incompatibility is not sufficient to falsify the TSS model since the failure of a small proportion of individuals to have reached a steady-state at overnight fasting is sufficient to explain the weak correlation. It is also likely that the assumption of fasting may not be true for the entire sample. Even if a small number of individuals do not comply with the overnight fasting instructions, a positive correlation can result and this possibility is extremely difficult to exclude in human data.

**Table 7:** Correlation and regression parameters of glucose-insulin relationship at steady and perturbed states.

|  | Steady-state(fasting) |  |  | Perturbed-state<br>(2 hours post glucose bolus) |  |  |
| --- | --- | --- | --- | --- | --- | --- |
| Parameter →<br>Data set ↓ | R-squared<br>(variance<br>explained) | p value | Slope (95%<br>CI) | R-squared<br>(variance<br>explained) | p value | Slope (95%<br>CI) |
| CRISIS<br>(N=522) | 0.0570 (5.7%) | <0.0001 | 0.1178<br>(0.0765 to<br>0.1591) | 0.3018<br>(30.18%) | <0.0001 | 1.2563<br>(1.0917 to<br>1.4209) |
| NHP-South Asian<br>(N=310) | 0.0172<br>(1.72%) | 0.021 | 0.1818<br>(0.0279 to<br>0.3356) | 0.344 (34.4%) | <0.0001 | 1.7347<br>(1.4661 to<br>2.0033) |
| NHP-European<br>(N=574) | 0.0548<br>(5.48%) | <0.0001 | 0.1988 (0.131<br>to 0.2666) | 0.2844<br>(28.44%) | <0.0001 | 0.8314<br>(0.7231 to<br>0.9397) |

The support of TSS model over CSS model is important because it accounts for the failure of impairment of insulin signalling to alter fasting glucose but increase only post load glucose.

## 7. Discussion

The five approaches examined above fail to support the classical belief about glucose insulin relationship. The insulin receptor knock-out experiments and insulin suppression or enhancement experiments converge to show that alteration in insulin levels or insulin sensitivity does not change the steady-state glucose levels. Evidence that it changes the shape of the glucose curve after food intake or glucose loading is more convincing in spite of some inconsistency across different experiments. Typically return to the steady-state is delayed by impaired insulin signalling but the steady-state glucose level remains unchanged.

Convergence of experiments using other means of causing specific alterations in insulin action strengthens the inference.

A number of mathematical models attempt to capture the dynamics of glucose homeostasis. A good model should be able to explain all the empirical results summed up here namely the inability of insulin receptor knockouts, insulin suppression and insulin enhancement experiments to alter steady state glucose levels; the difference in the regression correlation parameters between insulin and glucose in the steady versus perturbed state; the extremely weak correlation between fasting glucose and fasting insulin, but very strong correlation between HOMA-IR and HOMA-β; the hyperinsulinemic-normoglycemic prediabetic state and the phenomenon of impaired glucose tolerance but normal fasting glucose. Reviewing models of glucose homeostasis is beyond the scope of this paper, but we outline here what a good model of glucose homeostasis needs to explain. In our observation, all existing models explain only some of the empirical findings. We suggest here that this inability is because of a questionable common baseline assumption of all models that insulin signalling determines the glucose level in the fasting as well as post feeding conditions. It should be possible to construct such a model, if we realize that insulin affects glucose only in the post feeding but not in fasting conditions.

It is difficult to defend the classical assumptions about glucose-insulin relationship against the multiple convergent lines of evidence. Although results of these experiments have been there in the published literature for about two decades, these results were mostly explained away giving different excuses for different sets of experiments. The possible lines of defence would include difference between homeostatic mechanisms in rodents and humans or the possibility of non-linear nature of glucose-insulin relationship. The evidence reviewed here comes from rodents as well as humans and the glucose insulin scatters do not show any clear indication of non-linearity. Further it would be prudent to avoid making inferences based on dietary or other complex interventions since they can have multiple mechanisms of action.

Specific genetic or molecular interventions are more revealing with respect to the underlying mechanisms since we can be more confident about their specificity of action. Therefore our inference that insulin action does not influence fasting glucose levels is the most straightforward and parsimonious inference. Any other explanations will have to be supported by giving evidence for the assumptions made in those explanations.

The failure of experimental alteration in insulin signalling to alter steady-state glucose raises two distinct possibilities about fasting hyperglycaemia in T2D. One is that fasting hyperglycaemia in T2D is a result of processes independent of insulin signalling such as autonomic signalling or other insulin independent mechanisms. The sympathetic tone is known to be altered in metabolic syndrome (110) and increased sensitivity of liver to sympathetic signal is likely to be mainly responsible to fasting hyperglycaemia (111). The other possibility is that with impaired insulin signalling overnight fasting is not sufficient to reach a steady state, therefore fasting hyperglycaemia in T2D is a non-steady-state phenomenon in type 2 diabetes. The considerably weaker but still significant correlation between glucose and insulin in fasting as compared to post glucose load data suggests that both the factors are likely to be operational differentially in different individuals.

In either case certain fundamental concepts in our understanding of T2D need to be revised. First of all, the definition and measurement of insulin resistance using steady-state glucose and insulin levels needs to be questioned. Most commonly used indices of insulin resistance are based on the assumption that insulin signalling decides the fasting steady-state glucose levels, although non-equilibrium methods of assessing insulin resistance have been described (112). In the classical view other mechanisms of glucose regulation are assumed to be absent or non-significant. If increased sympathetic signalling increases liver glucose production, HOMA-IR will still account it as “insulin resistance”. The same is true about insulin resistance measured by hyperinsulinemic euglycemic clamp. The way insulin resistance is measured at the clinical level eliminates the chance of separately accounting for other mechanisms of glucose regulation. Even when experiments show that certain agents affect glucose dynamics independent of insulin action, they are typically labelled as “insulin sensitizing” agents (113). As a result, the belief that insulin is the only mechanism of glucose regulation relevant to T2D is artificially strengthened. There is a subtle circularity in the working definition of insulin resistance. Insulin resistance is blamed for the failure of normal or elevated levels of insulin to regulate glucose. In order to test this hypothesis, we should have an independent definition and measure of insulin resistance. Only then we can test whether and to what extent insulin resistance can alter glucose dynamics. However, clinically insulin resistance is measured by the inability of insulin to regulate glucose. Such a measure cannot be used to test the hypothesis that insulin resistance leads to the failure of insulin to regulate glucose. The unfalsifiability of the insulin resistance hypothesis arising out of this circularity has halted any attempts towards realistic assessments of the true causes of fasting hyperglycaemia in type 2 diabetes. In the molecular approach to induce insulin resistance, we have an independent definition and causality for insulin resistance and therefore such experiments are free from circularity of definition. The results of such experiments reviewed here are therefore more revealing and reliable. Since all of them converge to show that altering insulin signalling does not alter steady-state glucose levels, the insulin resistance and inadequate compensation hypothesis for steady-state hyperglycaemia stands clearly rejected.

The question can be turned upside down to examine whether steady-state glucose level determines steady-state insulin. If glucose is infused with a constant rate over a long time, insulin levels will come back to the baseline levels if glucose is not a determinant of fasting insulin. If it is, then insulin levels will stabilize at a new heightened steady-state level. Jetton et al. (2008) infused intra venous glucose (20% glucose w/v) continuously for 4 days in rats. Both glucose and insulin levels increased significantly after the infusion. However, later both glucose and insulin levels came back to normal even as the infusion continued. Increase in the concentration of the infused glucose (up to 35%) also yielded similar results (115). Thus, immediately on perturbation, glucose affected insulin levels, however after allowing sufficient time to regain steady state, the continued infusion of glucose had no significant effect on insulin levels. This demonstrates that even glucose does not hold a causal relationship with insulin in a steady-state whereas glucose level perturbation is certainly known to stimulate insulin response.

The interpretation of this phenomenon needs to be done at a broader philosophical level. We point out here with specific reference to homeostatic systems that the nature of causality in a perturbed-state can be qualitatively different from causality in steady state. There is a simple analogue to perturbed-state versus steady-state causality in one of the basic models of mathematical biology. In the classical model of logistic growth the intrinsic growth rate *r* decides the rate at which a population can change when away from the carrying capacity *K* (116). However, the carrying capacity itself may be independent of the growth rate. A non-zero positive *r* is required to reach the steady-state at *K* but *r* does not determine the steady-state level. It is a function of *K* alone. Reducing *r* leads to delay in achieving a steady state but the steady state remains at the same position. The evidence reviewed here indicates that insulin action is analogous to *r* of logistic model. It is required to reach a steady-state but it does not determine the location of the steady state.

The inability to distinguish between steady-state causality and perturbed-state causality may have substantially misled biomedical research at times, T2D certainly being an important example. This poses an important philosophical as well as methodological problem in experimental physiology. Many systems in physiology have homeostatic steady states and we use experimental approaches to reveal them. However, most experiments are perturbation experiments and we may be making the mistake of applying the demonstrated perturbed-state causality to understand steady-state systems. The apparent paradox can be resolved only by carefully designing and interpreting experiments. If a perturbation is momentary or transient, the results obtained would certainly reflect perturbed-state causality, but may not reflect steady-state causality. On the other hand, sustained perturbations held constant for sufficiently long to allow the system to regain a steady-state are necessary to establish steady-state causality. If upon sustainably altering a causal factor the effect variable returns to the same steady state, it reflects only perturbed-state and not steady-state causality. If, on the other hand, sustained alteration in the causal factor results into an altered steady state, it indicates steady-state causality.

Viewed from a slightly different and more generalized angle that goes beyond homeostatic systems, we can differentiate between two types of causalities. In driver causality the causal factor is necessary to reach a destination but does not decide the destination. In navigator causality the causal factor is crucial in determining the destination, but may not be sufficient to take the system there. The evidence reviewed above indicates that insulin is a driver but not a navigator of glucose homeostasis. A non-zero level of insulin is required for reaching a homeostatic steady state. In type 1 diabetes, the almost complete absence of insulin prevents glucose homeostasis. In type 2 diabetes there are non-zero insulin levels and therefore, a steady-state is possible, but insulin itself plays little role in deciding the steady-state glucose level. It is more likely that neuronal and other hormonal-metabolic factors affect the steady-state glucose in T2D.

Certain kinds of experimental interventions are unable to distinguish between driver versus navigator causality. Knocking out a driver or a navigator will disable the journey to the destination. Therefore, complete knockout of a cause may not distinguish between driver and navigator causality. On the other hand, experiments quantitatively altering the level of the causal factor while keeping it non-zero and observing the effect for sufficiently long duration, can help us differentiate between drivers and navigators. A sub-normal driver will delay the time to destination but will not change the destination. On the other hand, changing the navigator may or may not alter the time, but will alter the position of the destination. The history of insulin research is that early experiments such as total pancreatectomy demonstrated the necessary role of insulin in glucose homeostasis but the distinction between driver or navigator causality was not even conceptually perceived. So, it was assumed that insulin does both the roles. Although the absence of correlation between fasting glucose and insulin but good correlation after perturbation was noted as early as 1969 (117) but in the absence of conceptual differentiation between steady state and perturbed state causality, a clear interpretation did not emerge. Now in the presence of multiple experiments showing the precise role of insulin, we need to revive our concepts of causality. At a broader scale the insulin example warrants care in making inferences in experimental physiology, in the absence of which our understanding of the physiology of homeostatic systems can be seriously flawed.

## 8. Acknowledgements

We would like to thank Prof. Raj Bhopal, University of Edinburgh for providing clinical data from the Newcastle Heart Project. We would also like to thank Prof. Chittaranjan Yajnik, KEM Hospital, Pune for providing the clinical data from the CRISIS study. We are immensely grateful for the critical and insightful comments they provided on the earlier version of the manuscript. We would also like to thank Geetanjali Nerurkar for her help during the animal experiments.

## Supplementary information 1: Methods used for the meta-analyses

Four meta-analyses were performed in this study:

1. Insulin receptor knockout
2. Insulin degrading enzyme
3. Insulin suppression by diazoxide
4. Insulin suppression by octreotide

Methods used for the four meta-analyses:

**Keywords:** The keywords used in the meta analyses have been given in the table 1 of the main manuscript.

**Data-bases:** We have used the PubMed/MEDLINE database (and not the data bases which report clinical trials data) since the experiments we were searching for are predominantly experiments in basic research in life sciences as opposed to clinical studies. Majority of the studies which were searching for were rodent studies and not human studies.

**Timeline for inclusion of papers in the search:** The first search was performed in August 2017 and the papers until 31^st^ July 2017 were included in the primary search.

**Inclusion and exclusion criteria:** Given in the table 2 of the main paper

**Details of the papers:** Tables 1,2,3,4 below

**Methods of data extraction:** Data was extracted from the shortlisted papers using the software WebPlotDigitizer (Author: Ankit Rohatgi Website: https://automeris.io/WebPlotDigitizer, Version: 4.1, January, 2018, E-Mail, Location: Austin, Texas, USA)

**Principal summary measures:** The data extracted from each shortlisted paper was the difference of means of blood/plasma glucose levels between the ‘control’ and the ‘treated’ along with the 95% confidence intervals.

**Methods of handling data and combining results of studies:** These differences in the means between the control and treated from all the respective shortlisted papers were compiled. These differences were compared across different timepoints using the non-parametric chi-square test.

**Details of the papers shortlisted for the four meta-analyses:**

**Table 1:**
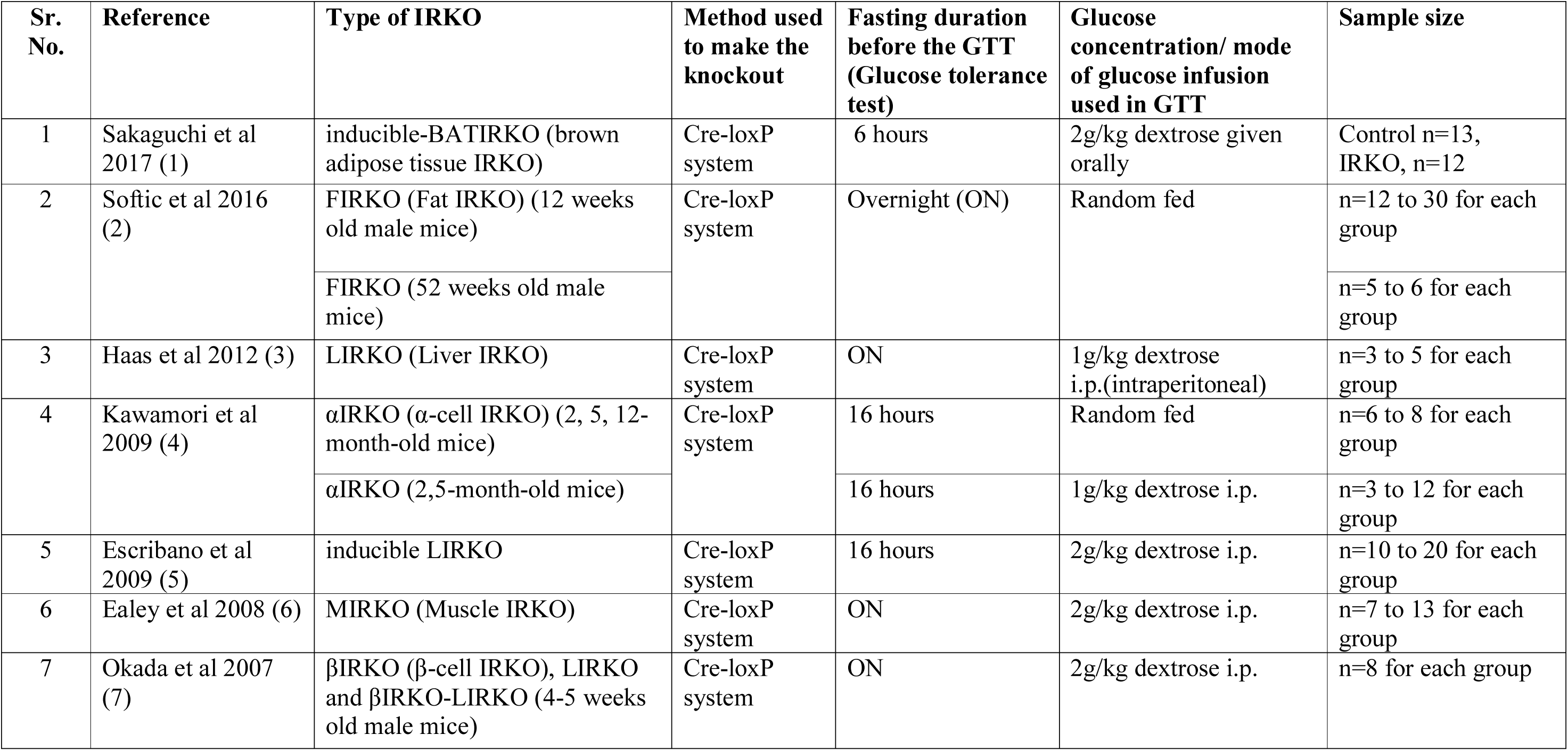
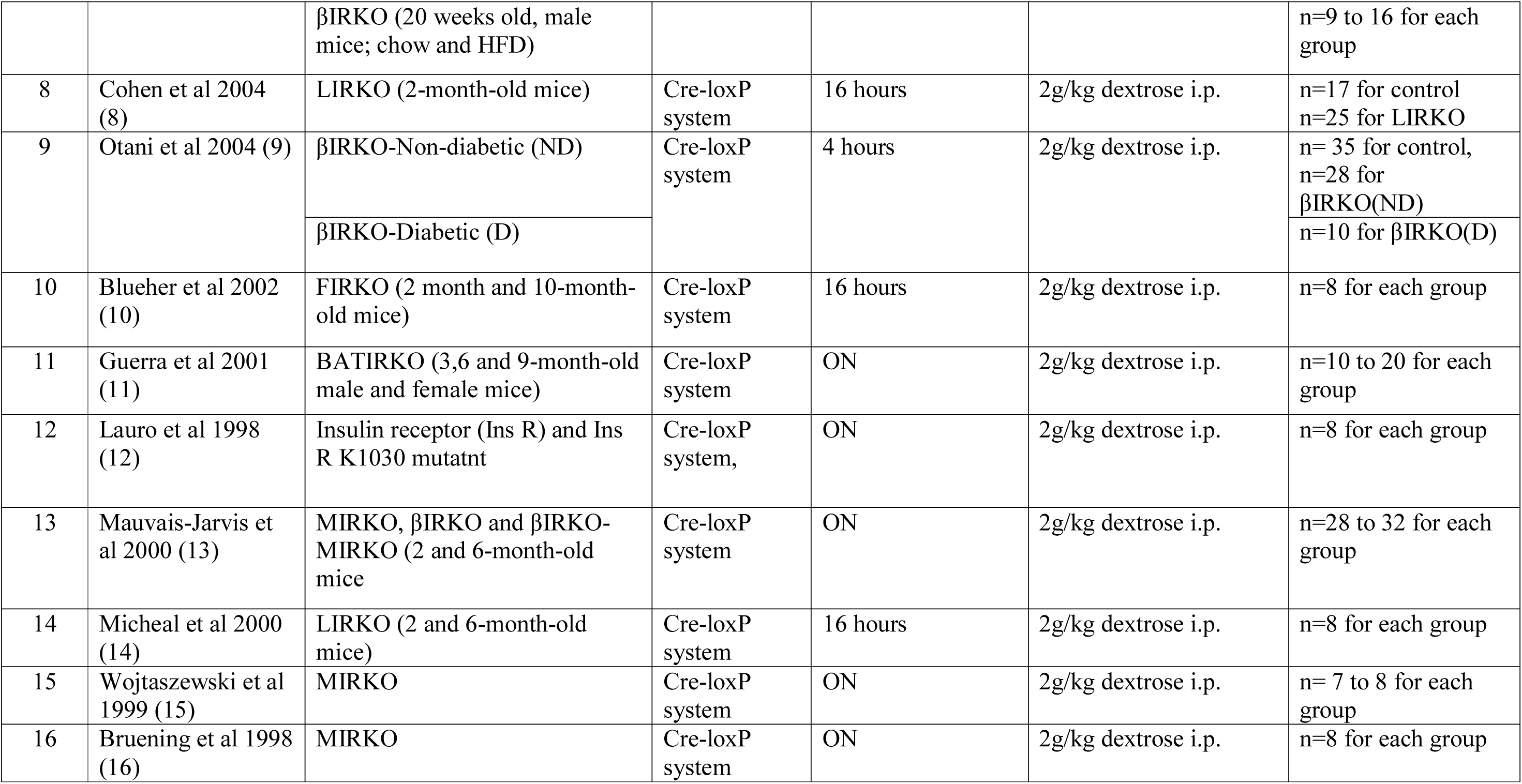
Details of the 16 papers used in the Insulin Receptor Knock-Out (IRKO) meta-analysis. All of these studies were carried out on rodent models.

**Table 2:** Details of the 6 papers used in the Insulin Degrading Enzyme (IDE) inhibition meta-analysis. All studies were carried out on rodent models.

| Sr. No. | Reference | Method used to inhibit IDE | Fasting duration before the GTT | Glucose concentration/ mode of glucose infusion used in GTT | Sample size |
| --- | --- | --- | --- | --- | --- |
| 1 | Villa Perez et al 2018 (17) | Liver specific IDE knockout | 16 hours | 2g/kg dextrose given i.p. | n= 9 to 13 for each group |
| 2 | Deprez-Poulain et al 2018 (18) | Inhibition of catalytic site of IDE using the inhibitor BDM44768 | 6 hours | 1.5g/kg glucose for IPGTT and 2 or 3g/kg glucose for OGTT | n= 4 to 7 for each group |
| 3 | Durham et al 2015 (19) | Inhibition of IDE using an N-terminal exosite (NTE) | ON | 2g/kg dextrose given orally | n=6 for each group |
| 4 | Maianti et al 2014 (20) | Inhibition of IDE using a non-catalytic site binding inhibitor | 14 hours | 1.5g/kg glucose for IPGTT and 3g/kg glucose for OGTT | n=5 to 7 for each group |
| 5 | Abdul Hay et al 2011 (21) | IDE-KO created by Cre-lox recombination | 6 to 9 hours | 1g/kg dextrose given i.p. | n=10 to 12 for each group |
| 6 | Farris et al 2003 (22) | IDE<sup>-/-</sup> mice created by gene trapping method | ON | 2g/kg dextrose given i.p. | n=6 (IDE<sup>-/-</sup>)<br>n=4 (Control) |

**Table 3:**
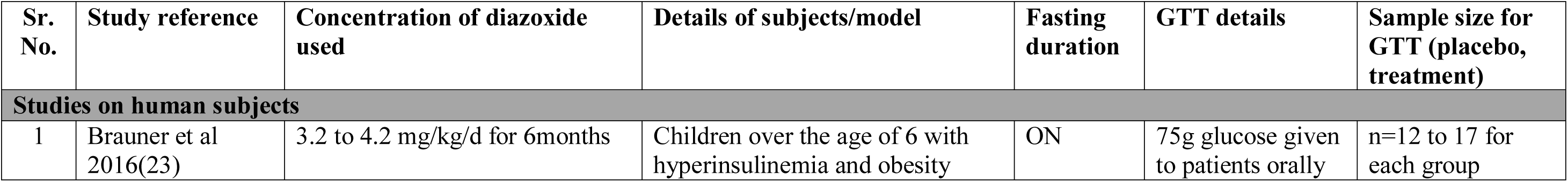
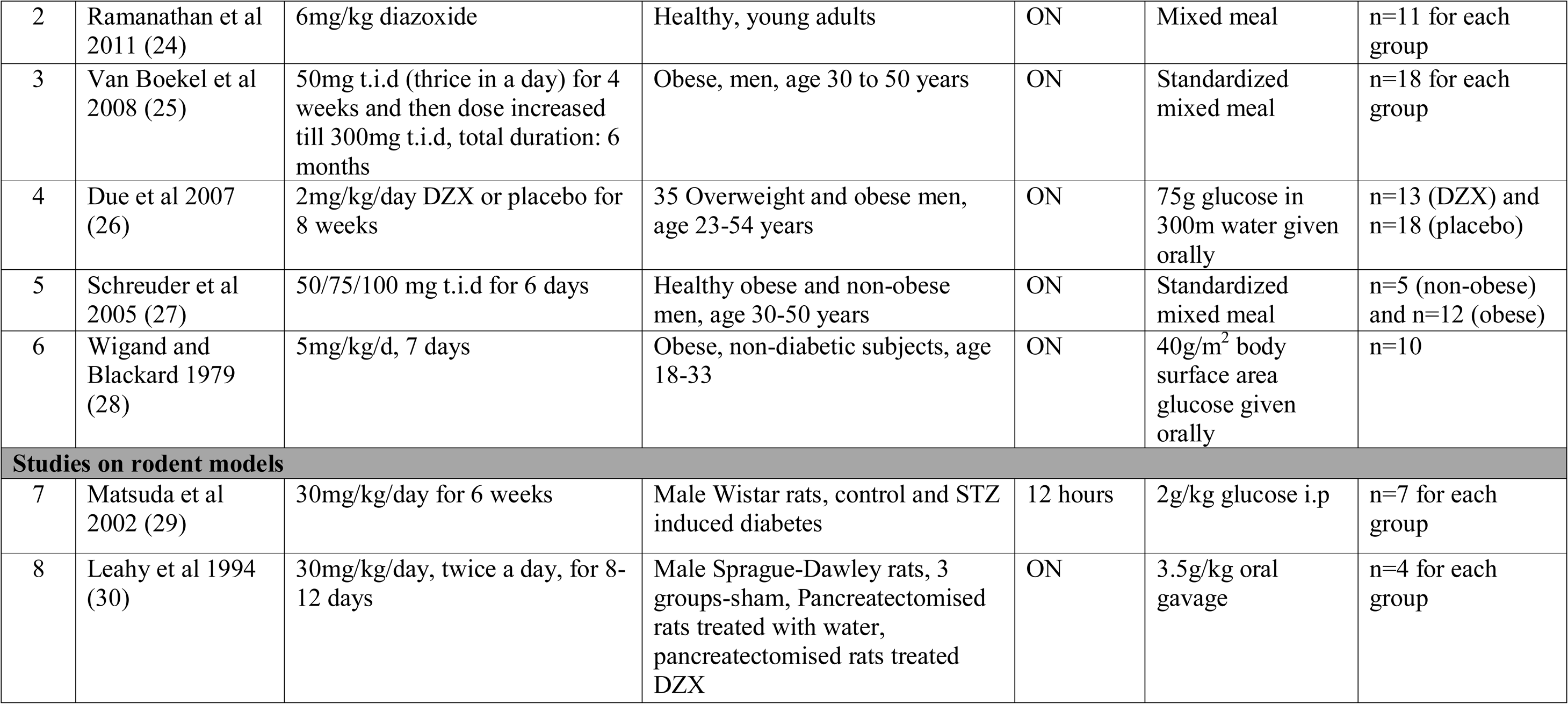
Details of the 8 papers used in the Diazoxide (DZX) meta-analysis.

| Sr. No. | Study reference | Concentration of diazoxide used | Details of subjects/model | Fasting duration | GTT details | Sample size for GTT (placebo, treatment) |
| --- | --- | --- | --- | --- | --- | --- |
| **Studies on human subjects** |  |  |  |  |  |  |
| 1 | Brauner et al 2016(23) | 3.2 to 4.2 mg/kg/d for 6months | Children over the age of 6 with hyperinsulinemia and obesity | ON | 75g glucose given to patients orally | n=12 to 17 for each group |
| 2 | Ramanathan et al 2011 (24) | 6mg/kg diazoxide | Healthy, young adults | ON | Mixed meal | n=11 for each group |
| 3 | Van Boekel et al 2008 (25) | 50mg t.i.d (thrice in a day) for 4 weeks and then dose increased till 300mg t.i.d, total duration: 6 months | Obese, men, age 30 to 50 years | ON | Standardized mixed meal | n=18 for each group |
| 4 | Due et al 2007 (26) | 2mg/kg/day DZX or placebo for 8 weeks | 35 Overweight and obese men, age 23-54 years | ON | 75g glucose in 300ml water given orally | n=13 (DZX) and n=18 (placebo) |
| 5 | Schreuder et al 2005 (27) | 50/75/100 mg t.i.d for 6 days | Healthy obese and non-obese men, age 30-50 years | ON | Standardized mixed meal | n=5 (non-obese) and n=12 (obese) |
| 6 | Wigand and Blackard 1979 (28) | 5mg/kg/d, 7 days | Obese, non-diabetic subjects, age 18-33 | ON | 40g/m<sup>2</sup> body surface area glucose given orally | n=10 |
| **Studies on rodent models** |  |  |  |  |  |  |
| 7 | Matsuda et al 2002 (29) | 30mg/kg/day for 6 weeks | Male Wistar rats, control and STZ induced diabetes | 12 hours | 2g/kg glucose i.p | n=7 for each group |
| 8 | Leahy et al 1994 (30) | 30mg/kg/day, twice a day, for 8-12 days | Male Sprague-Dawley rats, 3 groups-sham, Pancreatectomised rats treated with water, pancreatectomised rats treated DZX | ON | 3.5g/kg oral gavage | n=4 for each group |

**Table 4:** Details of the 10 papers used in the Octreotide (OCT) meta-analysis. All the papers included studies on human subjects.

| Sr. No. | Study reference | Concentration of octreotide used | Details of subjects/model | Fasting duration | GTT details | Sample size for GTT (placebo, treatment) |
| --- | --- | --- | --- | --- | --- | --- |
| 1 | Madsen et al 2011 (31) | <u>Somatostatin analog (SA) alone:</u><br>Octreotide 10-30mg/4weeks OR<br>Lanreotide 80mg/4weeks<br><u>Co-treatment:</u><br>Octreotide 6.7-20mg/4weeks OR<br>Lanreotide 24-60mg/4weeks<br>AND<br>Pegvisomat 30-60mg/4weeks | 18 Acromegalic patients (age 54±3 years) | ON | 75g glucose given orally | n=6 SA only<br>n=12 SA+P |
| 2 | Breckenbridge et al 2007 (32) | Octreotide (30ng/kg.min) with GH +<br>1. Saline (treatment for our purpose)<br>2. Insulin (control for our purpose) | 14 Healthy adults (BMI 23 ± 2.9; Age 29 ± 5 years) | ON | 22.5µmol/kg | n=8 male<br>n=6 female |
| 3 | Ronchi et al 2004 (33) | 1. Lanreotide (Slow Release)-30mg im injection every 14 days for 19±16 months<br>2. Octreotide (Long Acting Release)-20mg im injection every 28 days for 21±10 months | 10 acromegalic patients (6 men and 4 women; age 46±16 years; BMI 29±5) | ON | not mentioned | n=6 male<br>n=4 female |
| 4 | Parkinson et al 2002 (34) | 1. Octreotide (50µg sc t.i.d) for 7 days<br>2. Pegvisomant (20mg/day sc) for 7 days | 6 healthy, male volunteers (age 21-63 years), studied on 3 separate occasions | ON | 75g glucose given orally | n=6 male |
| 5 | Giustina et al 1991 (35) | T2D patients received either of the four treatments<br>a. Insulin 0.1U/kg<br>b. Octreotide 25µg<br>c. Oct 50µg<br>d. Oct 100µg with insulin | 8 overweight/obese T2D patients (age $53.4 \pm 4.2$ years) | ON | Mixed meal | n=8 (7 female and 1 male) |
| 6 | Candrina and Giustina 1998 (36) | Type 2 diabetes patients received 0.5U/kg/day divided into 2 subcutaneous injections | 5 T2D patients (age $56 \pm 4$ years) duration of diabetes ranged from 13 to 25 years | ON | 300 kcal breakfast | n=5 (3 male and 2 female) |
| 7 | Williams et al 1988 (37) | Type 2 diabetes patients received 50µg OCT, thrice a day, subcutaneously, for 3 days | 7 T2D patients (age 51-73 years, mean 67 years), duration of diabetes ranged from (6 months to 4 years, mean 2.5 years) | ON | Standardised breakfast | n=7 (4 males and 3 females) |
| 8 | Johnston et al 1986 (38) | 50 µg OCT administered subcutaneously, twice a day in diabetic and nondiabetic patients | 6 normal and 5 type 2 diabetic subjects (age range not given) | ON | Standardised meals | n=6 (normal, male)<br>n=5 (T2D) |
| 9 | Davies et al 1986 (39) | 50 µg OCT administered subcutaneously, twice a day in diabetic patients | 5 T2D patients (mean age 49 years; duration of diabetes ranged from 3 to 10 years, with a mean of 6 years) | ON | Standardised breakfast | n=5 (male, diabetic) |
| 10 | Williams et al 1986 (40) | 5-100µg OCT administered subcutaneously in T2D patients twice a day<br>Concentrations used:<br>50 µg in normal<br>5,100 µg in T2D | 5 normal and 5 T2D patients (age 50 to 65 years) | ON | Standardised breakfast | n=5 (normal)<br>n=5 (T2D) |

## Supplementary information 2: A generalized CSS model to make predictions testable in population data

A number of models of glucose regulation exist in literature. We use a simple model assuming the following. The plasma glucose level *G* increases by two processes namely absorption from gut and glucose production by the liver. We assume the gut absorption *Gt* to be independent of standing plasma glucose as well as insulin, whereas liver glucose production has a maximum rate *L* which has two feedback inhibitors namely direct feedback inhibition by glucose and that by standing plasma insulin which depends upon the insulin sensitivity of liver. Glucose clearance has two mechanisms namely insulin independent and insulin dependent. The plasma insulin *I* is a balance between insulin release by pancreatic beta cells, the rate being a function of plasma glucose and a rate of insulin degradation which is directly proportional to standing plasma insulin level. We assume all relationships to be linear and use the model framework of Chawla et al 2018 (1).

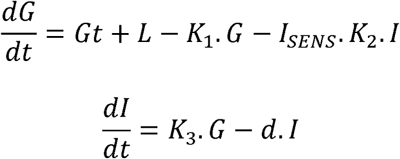

Where *K_1_* is a rate constant for glucose uptake by tissues as well as direct feedback inhibition of liver glucose production, *K_2_* a rate constant for insulin mediated inhibition of liver glucose production as well as insulin mediated glucose uptake, both of which are assumed to be a function of insulin sensitivity *I_SENS_* which is assumed to be unity normally and decreases with insulin resistance. *K_3_* is the rate constant for glucose stimulated insulin secretion and *d* the rate of insulin clearance.

We use simulations with normally distributed errors to study how the correlation between plasma glucose and insulin is affected by the parameters as well as by the standard deviation of errors. We use the errors additively or multiplicatively. For simulations using additive errors, we add normally distributed error terms *e_1_* and *e_2_* to both the equations.

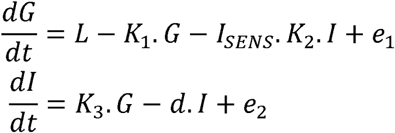

For simulations using multiplicative error, we give normal distributions to *K_1_, K_2_, K_3_* and *I_SENS_*. Realistic ranges for the parameters are taken from Chawla et al 2018(1).

Simulations show that in a additive error model, as long as the parameters of glucose insulin relationship are the same, the regression correlation parameters for glucose insulin relationship are not significantly different during fasting steady state (*Gt=0*) and at any time post meal (*Gt > 0*). The only difference is in the range of glucose and insulin distribution (figure 1)

**Figure 1:**
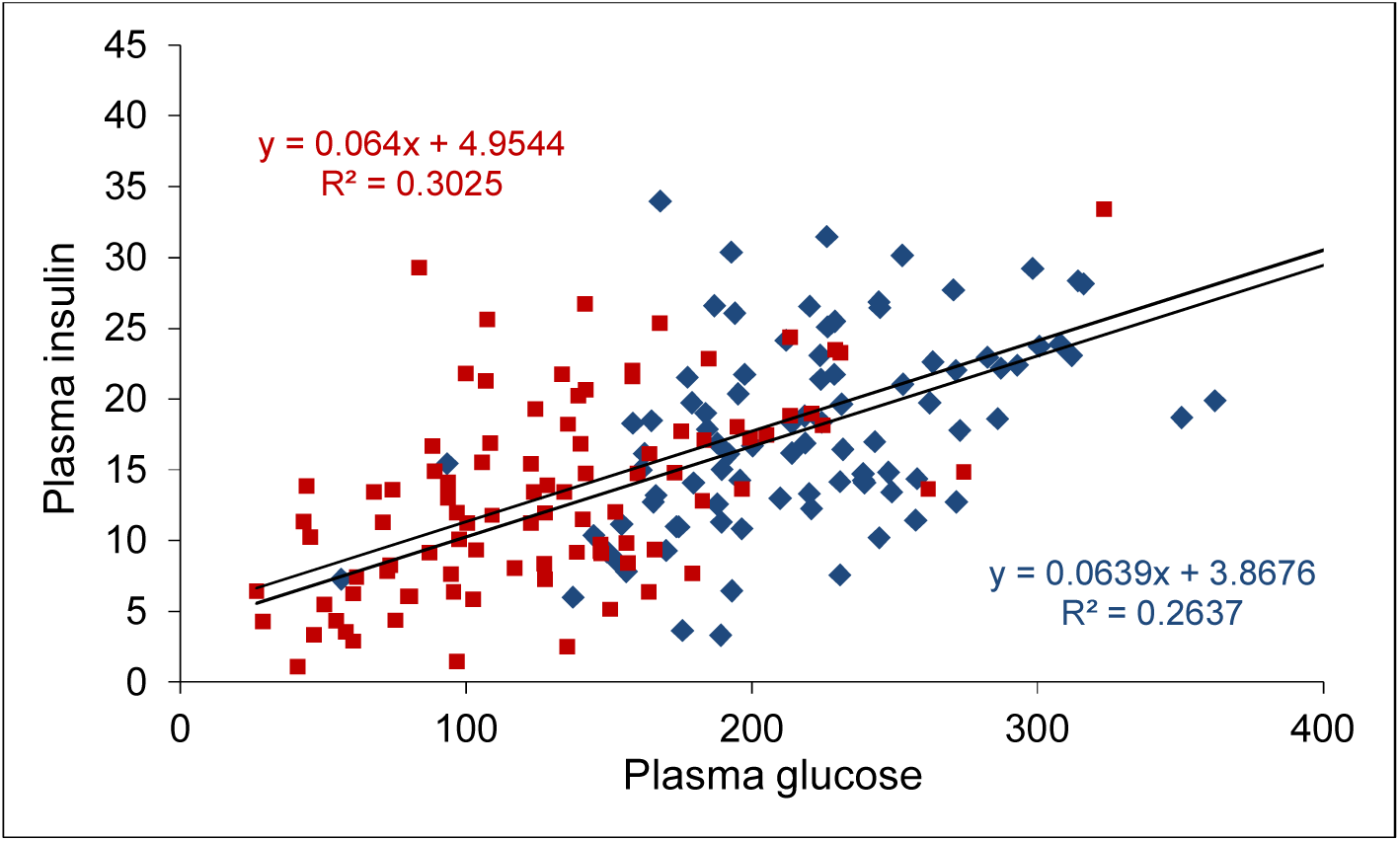
The glucose insulin scatter in a fasting steady state (red squares) and in a post meal arbitrary but constant time interval (blue diamonds) in an additive error model. A sample result is shown in which *K_1_*=0.1, *K_2_*=0.9, *I_SENS_* is randomized between 0.1 and 1 and *K_3_*=0.015 and *d*=0.15. The error standard deviations are 15 and 1 respectively.

In simulations with multiplicative errors, the post meal glucose insulin correlation was always weaker than the fasting steady state correlation (figure 2). This is the likely result of the errors growing in proportion to larger values of glucose and insulin, and also due to an additional variable, gut absorption being incorporated in the model.

The results were not sensitive to parameter changes as long as G and I were positive. We can confidently make a generalization that as long as the model parameters remain the same, the glucose insulin correlation in steady state is stronger or equal to the post meal correlation.

Logically and intuitively sound, this generalization is unlikely to be specific to a particular form of equations based on the assumptions of the CSS class of models.

**Figure 2:**
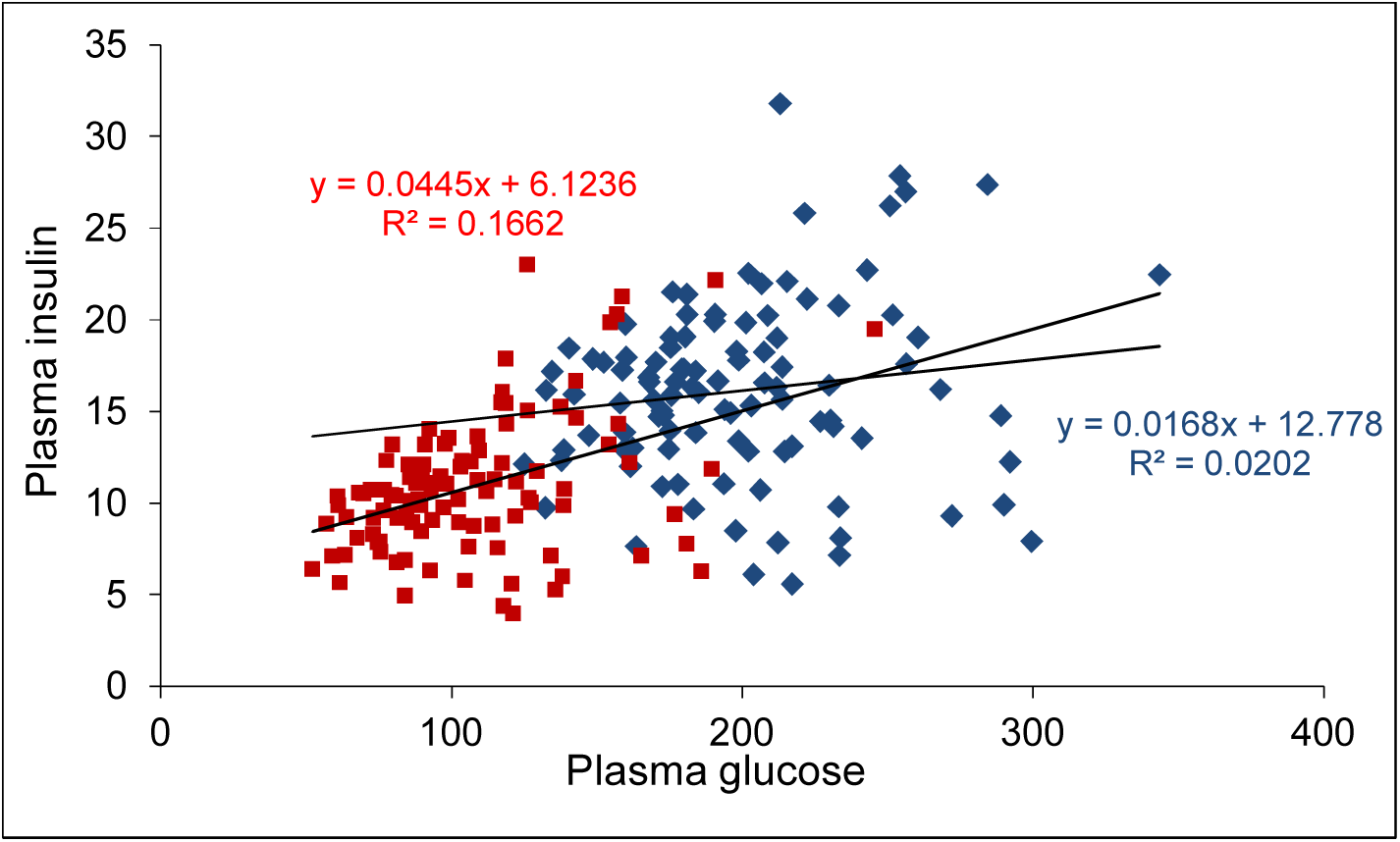
The glucose insulin scatter in a fasting steady state (red squares) and at a post meal arbitrary but constant time point (blue diamonds) in a multiplicative error model. A sample result is shown in which the mean (standard deviations) of the parameters were *K_1_*=0.1 (0.02), *K_2_*=0.9 (0.5), *I_SENS_* is randomized between 0.1 and 1 *K_3_*=0.0015 (0.0002) and *d* =0.15 (0.005). In all the simulations the correlation coefficient and regression slopes of the post meal scatters were less than or equal to the corresponding fasting parameters. This contrasts the epidemiological patterns in which the fasting correlations are substantially weaker than the post meal correlations (main text Table 7 figure 10).

The simulation results contrast with real life data in which the steady state correlation and regression slope between glucose and insulin is observed to be substantially weaker than the post meal relations at any point in time. This indicates that the parameters of glucose insulin relationship in steady state are substantially different from the post meal parameters, or glucose insulin relationship in steady state is qualitatively different from that in the perturbed state.

## Supplementary 1283 information 3

1. PRISMA 2009 Checklist for the Insulin Receptor Knockout meta-analysis.

| Section/topic | # | Checklist item | Reported on page # |
| --- | --- | --- | --- |
| **TITLE** |  |  |  |
| Title | 1 | Identify the report as a systematic review, meta-analysis, or both. | Not applicable (NA) since meta-analyses is one of the methods used to address the question and not the only method. |
| **ABSTRACT** |  |  |  |
| Structured summary | 2 | Provide a structured summary including, as applicable: background; objectives; data sources; study eligibility criteria, participants, and interventions; study appraisal and synthesis methods; results; limitations; conclusions and implications of key findings; systematic review registration number. | 2 |
| **INTRODUCTION** |  |  |  |
| Rationale | 3 | Describe the rationale for the review in the context of what is already known. | Page 5, Line 50 |
| Objectives | 4 | Provide an explicit statement of questions being addressed with reference to participants, interventions, comparisons, outcomes, and study design (PICOS). | Page 6 |
| **METHODS** |  |  |  |
| Protocol and registration | 5 | Indicate if a review protocol exists, if and where it can be accessed (e.g., Web address), and, if available, provide registration information including registration number. | NA |
| Eligibility criteria | 6 | Specify study characteristics (e.g., PICOS, length of follow-up) and report characteristics (e.g., years considered, language, publication status) used as criteria for eligibility, giving rationale. | NA |
| Information sources | 7 | Describe all information sources (e.g., databases with dates of coverage, contact with study authors to identify additional studies) in the search and date last searched. | Page 7: Table 1;<br>Page 51: Supplementary information 1 |
| Search | 8 | Present full electronic search strategy for at least one database, including any limits used, such that it could be repeated. | Page 7: Table 1;<br>Page 51-52: Supplementary information 1 |
| Study selection | 9 | State the process for selecting studies (i.e., screening, eligibility, included in systematic review, and, if applicable, included in the meta-analysis). | Page 7: Table 1;<br>Page 51-52: Supplementary information 1 |
| Data collection process | 10 | Describe method of data extraction from reports (e.g., piloted forms, independently, in duplicate) and any processes for obtaining and confirming data from investigators. | Data extraction: Page 51: Supplementary information 1<br>No process for obtaining and confirming data from investigators, only published data used. |
| Data items | 11 | List and define all variables for which data were sought (e.g., PICOS, funding sources) and any assumptions and simplifications made. | Page 51-52: Supplementary information 1 |
| Risk of bias in individual studies | 12 | Describe methods used for assessing risk of bias of individual studies (including specification of whether this was done at the study or outcome level), and how this information is to be used in any data synthesis. | NA |
| Summary measures | 13 | State the principal summary measures (e.g., risk ratio, difference in means). | Difference in means with the 95% Confidence Intervals of the means. (Page 51-52: Supplementary information 1) |
| Synthesis of results | 14 | Describe the methods of handling data and combining results of studies, if done, including measures of consistency (e.g., $I^2$) for each meta-analysis. | NA |
| Risk of bias across studies | 15 | Specify any assessment of risk of bias that may affect the cumulative evidence (e.g., publication bias, selective reporting within studies). | NA |
| Additional analyses | 16 | Describe methods of additional analyses (e.g., sensitivity or subgroup analyses, meta-regression), if done, indicating which were pre-specified. | NA |
| **RESULTS** |  |  |  |
| Study selection | 17 | Give numbers of studies screened, assessed for eligibility, and included in the review, with reasons for exclusions at each stage, ideally with a flow diagram. | Page 7: Table 1; |
| Study characteristics | 18 | For each study, present characteristics for which data were extracted (e.g., study size, PICOS, follow-up period) and provide the citations. | Page 51-52: Supplementary information 1 |
| Risk of bias within studies | 19 | Present data on risk of bias of each study and, if available, any outcome level assessment (see item 12). | NA |
| Results of individual studies | 20 | For all outcomes considered (benefits or harms), present, for each study: (a) simple summary data for each intervention group (b) effect estimates and confidence intervals, ideally with a forest plot. | (a) Pages 51 to 52: Supplementary information 1<br>(b) Figures 1 to 4 in the paper<br>(c) Table 2 in the paper |
| Synthesis of results | 21 | Present results of each meta-analysis done, including confidence intervals and measures of consistency. | Pages 8-15 in the paper |
| Risk of bias across studies | 22 | Present results of any assessment of risk of bias across studies (see Item 15). | NA |
| Additional analysis | 23 | Give results of additional analyses, if done (e.g., sensitivity or subgroup analyses, meta-regression [see Item 16]). | NA |
| **DISCUSSION** |  |  |  |
| Summary of evidence | 24 | Summarize the main findings including the strength of evidence for each main outcome; consider their relevance to key groups (e.g., healthcare providers, users, and policy makers). | Pages 37-43 |
| Limitations | 25 | Discuss limitations at study and outcome level (e.g., risk of bias), and at review-level (e.g., incomplete retrieval of identified research, reporting bias). | Pages 37-43 |
| Conclusions | 26 | Provide a general interpretation of the results in the context of other evidence, and implications for future research. | Pages 37-43 |
| **FUNDING** |  |  |  |
| Funding | 27 | Describe sources of funding for the systematic review and other support (e.g., supply of data); role of funders for the systematic review. | Page 43 |

2. PRISMA 2009 Checklist for the Insulin Degrading Enzyme meta-analysis.

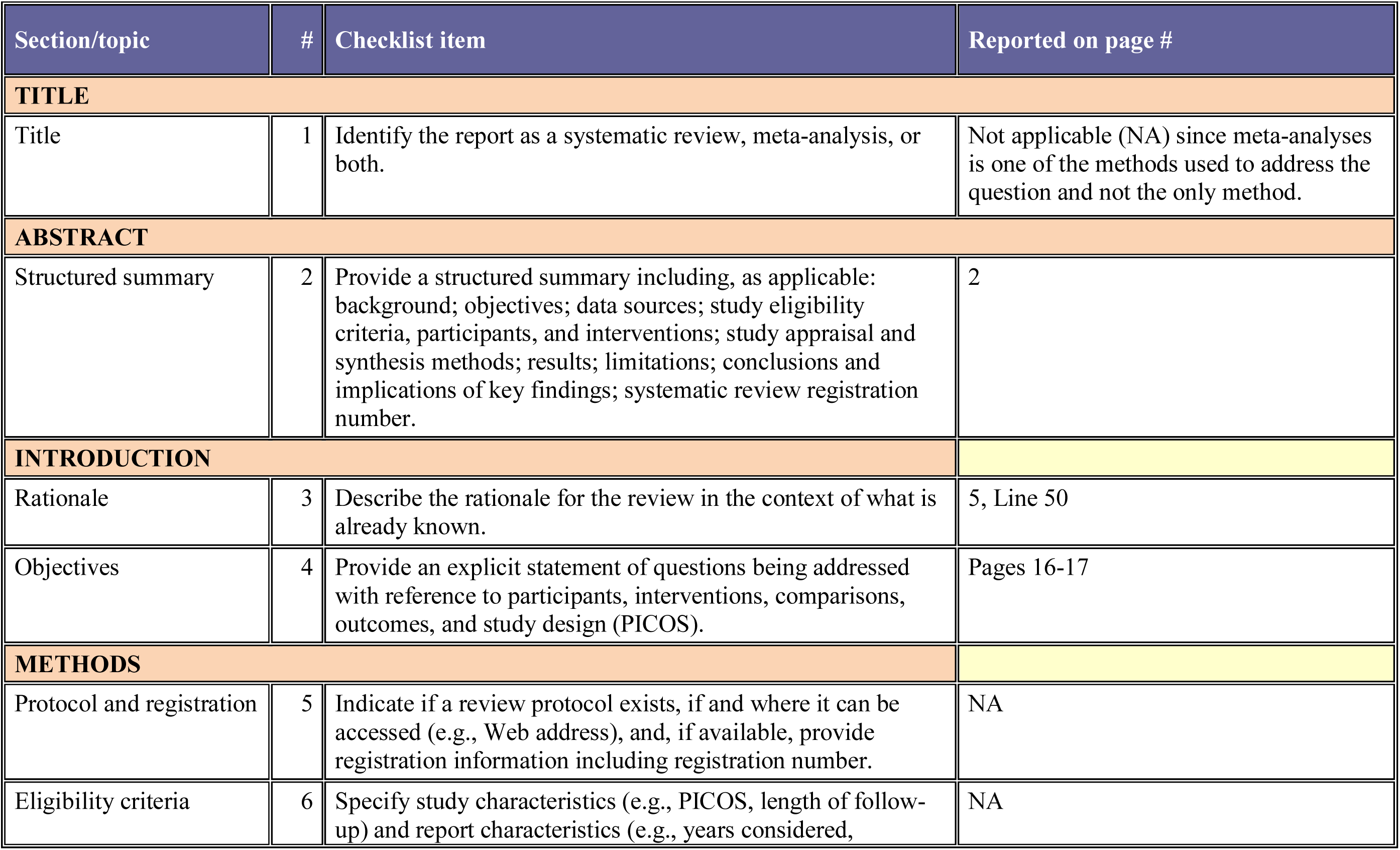
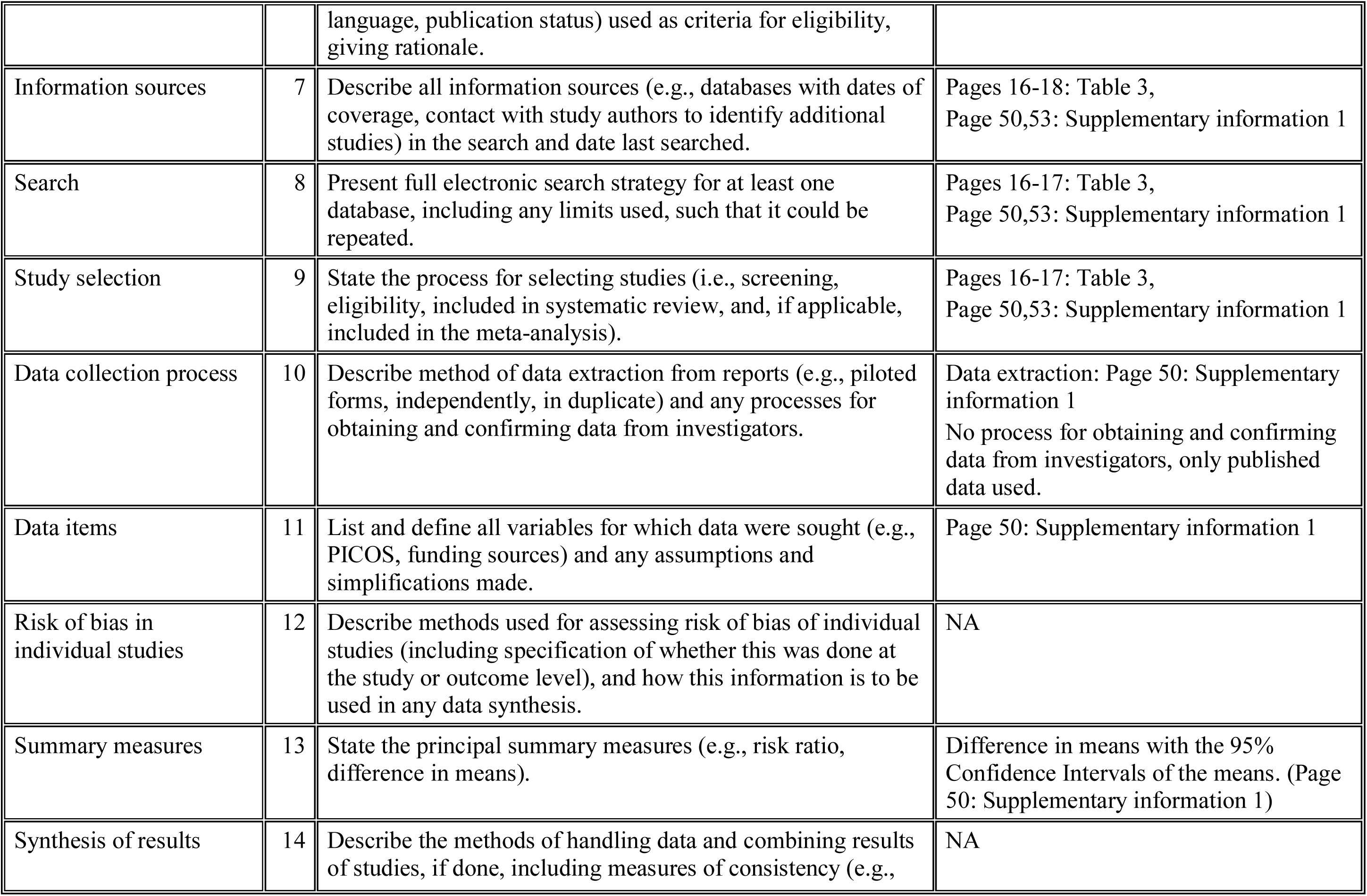
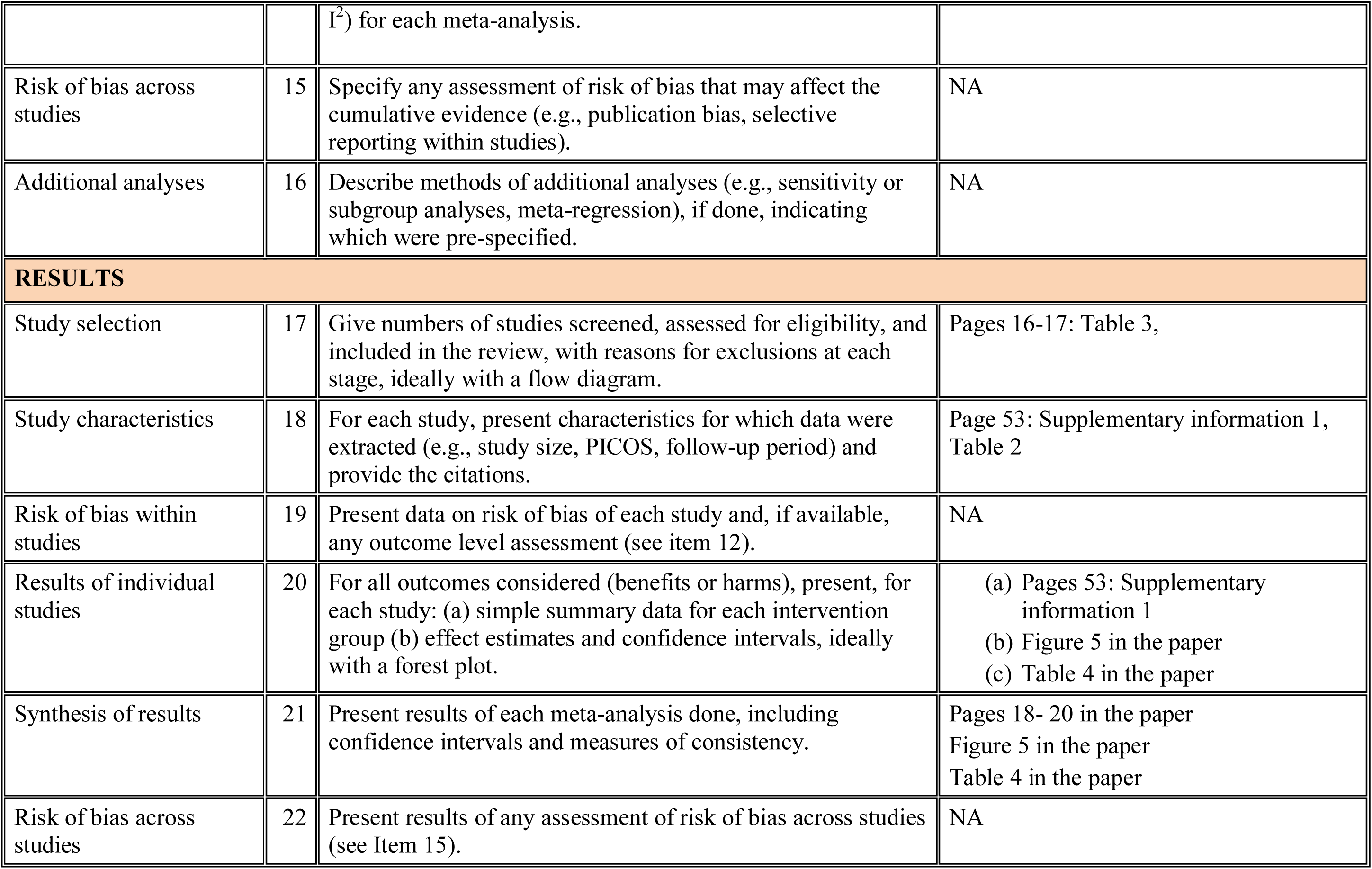
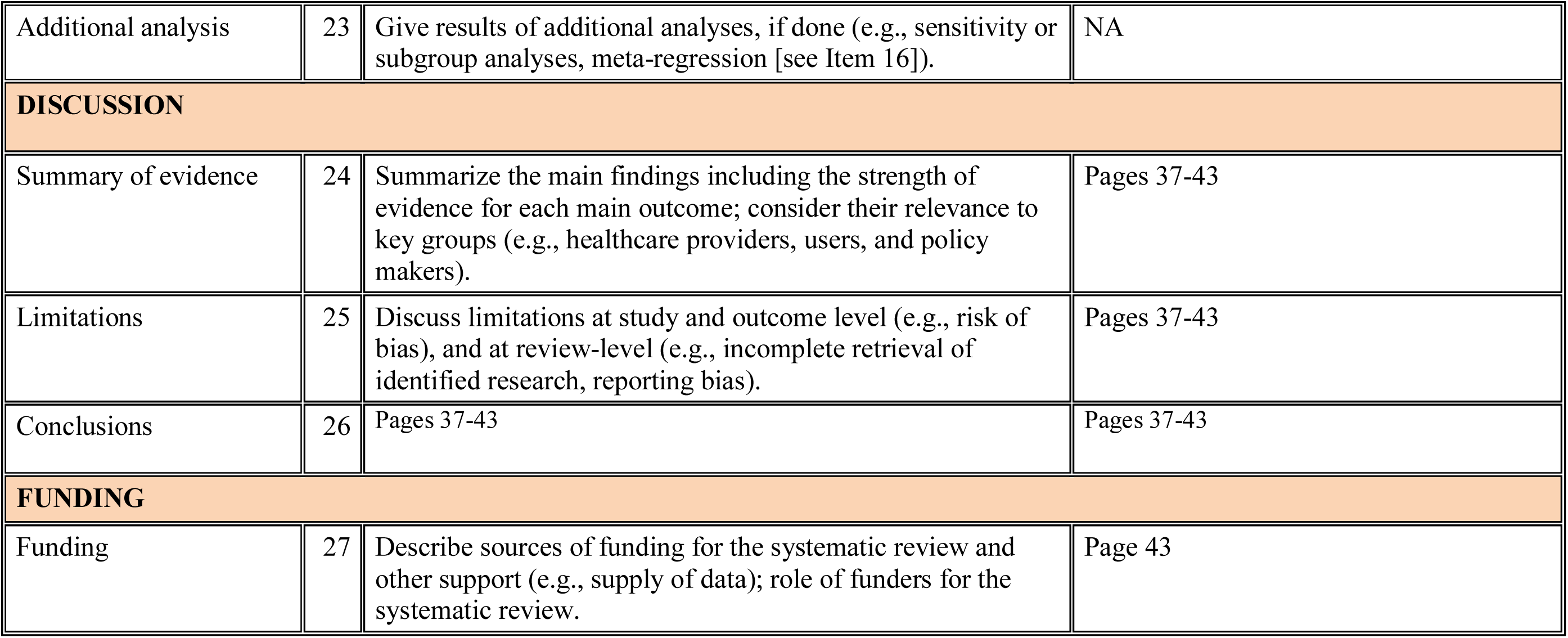

3. PRISMA 2009 Checklist for the Insulin Suppression by Diazoxide meta-analysis.

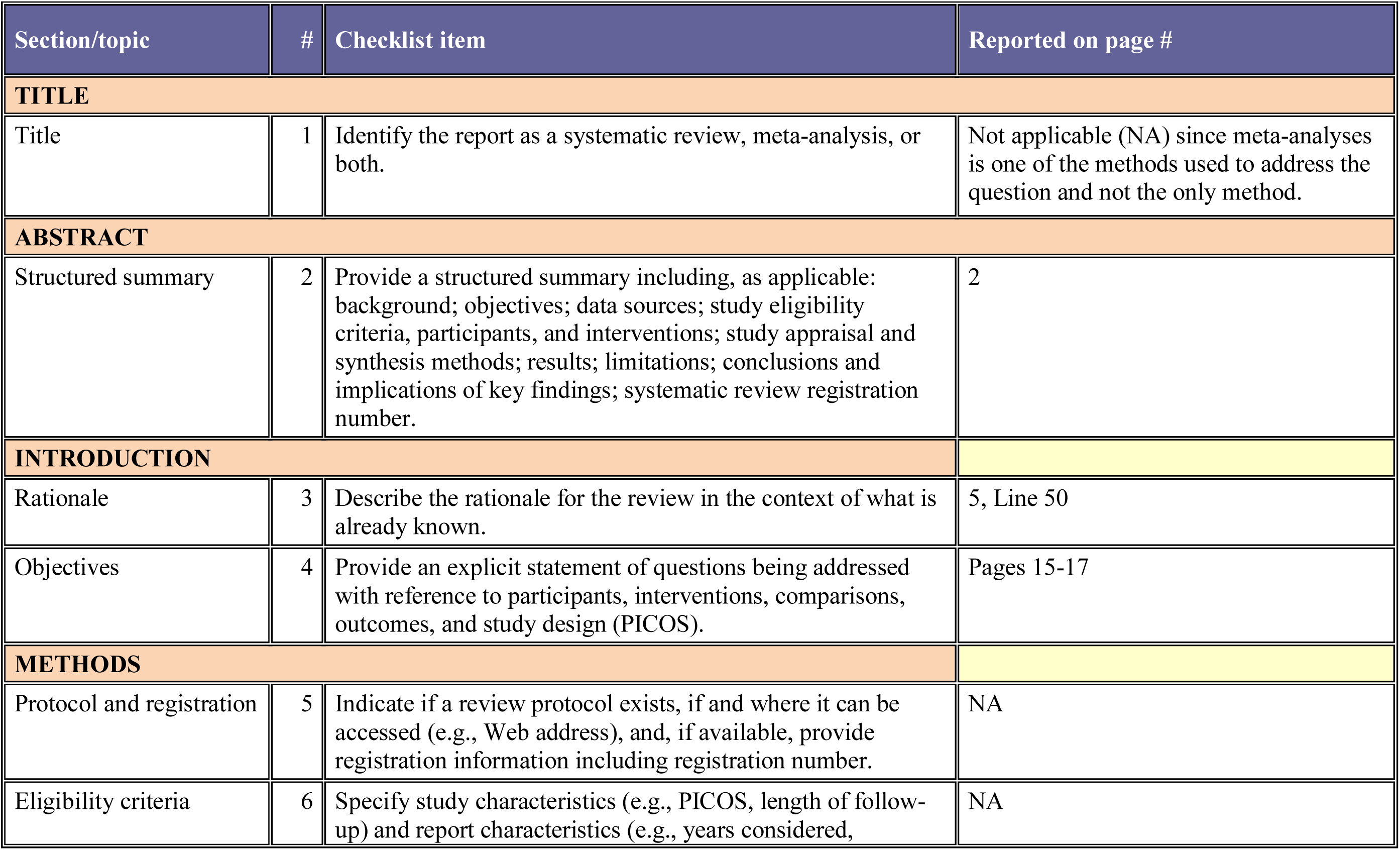
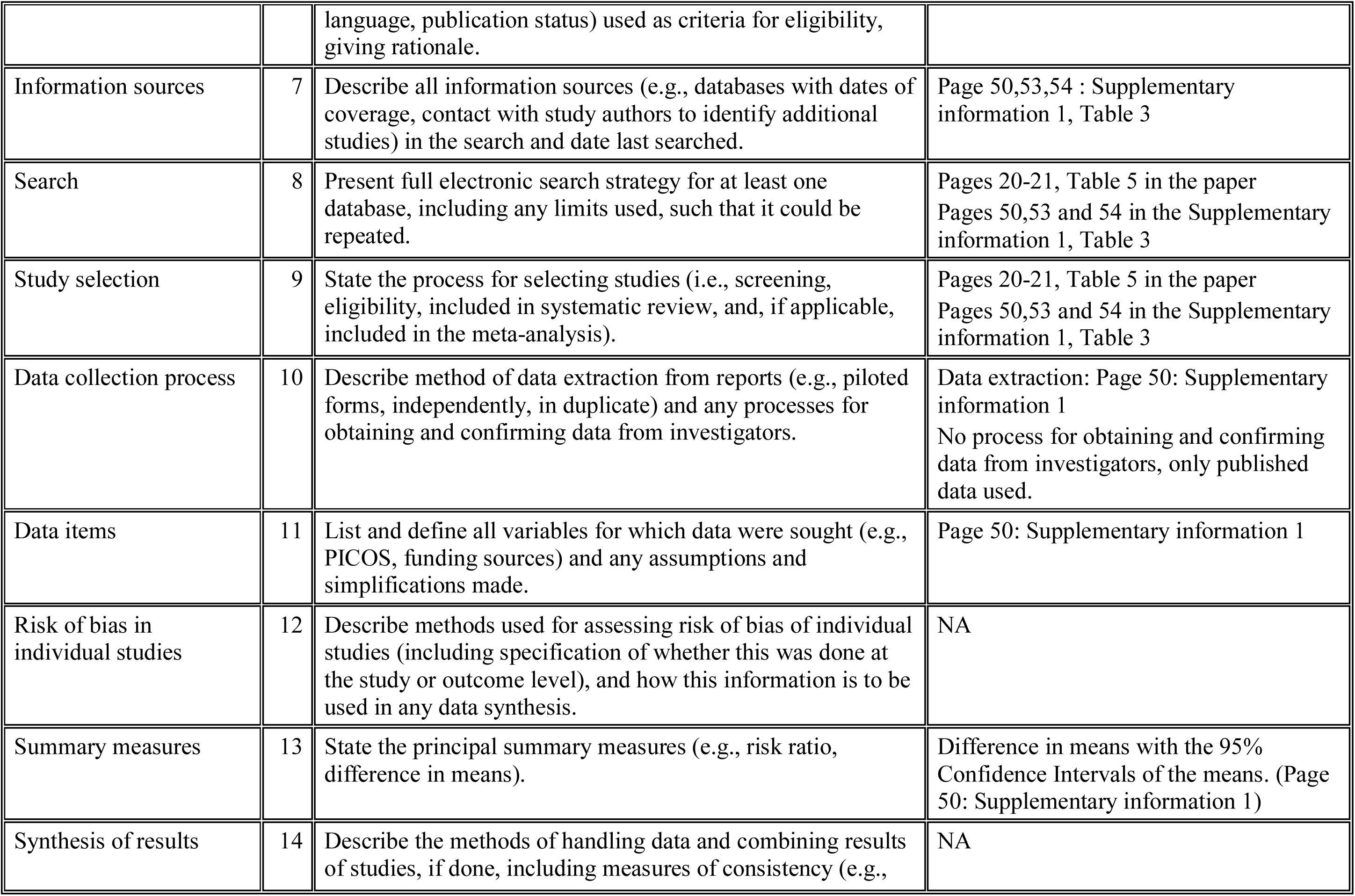
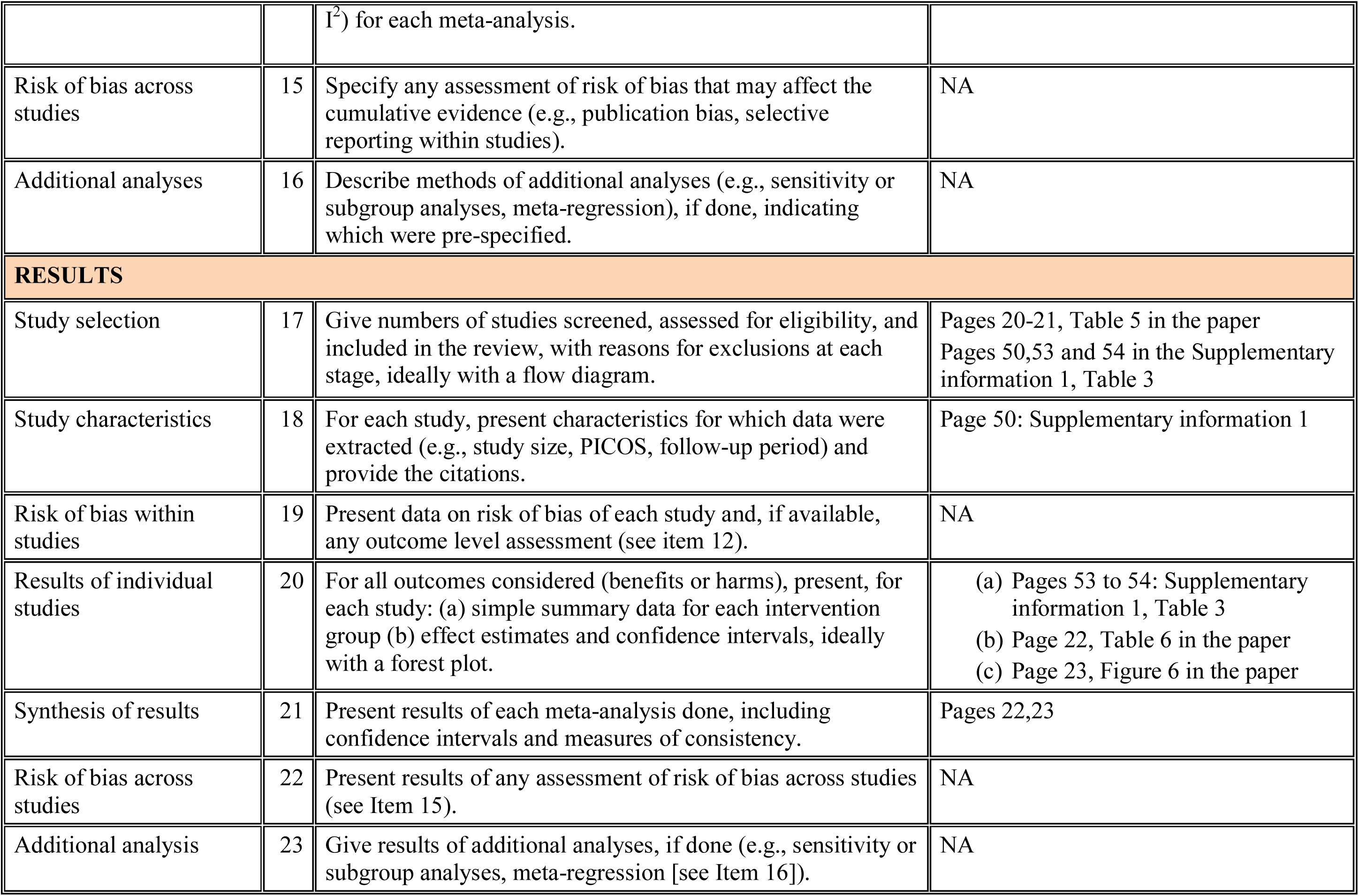
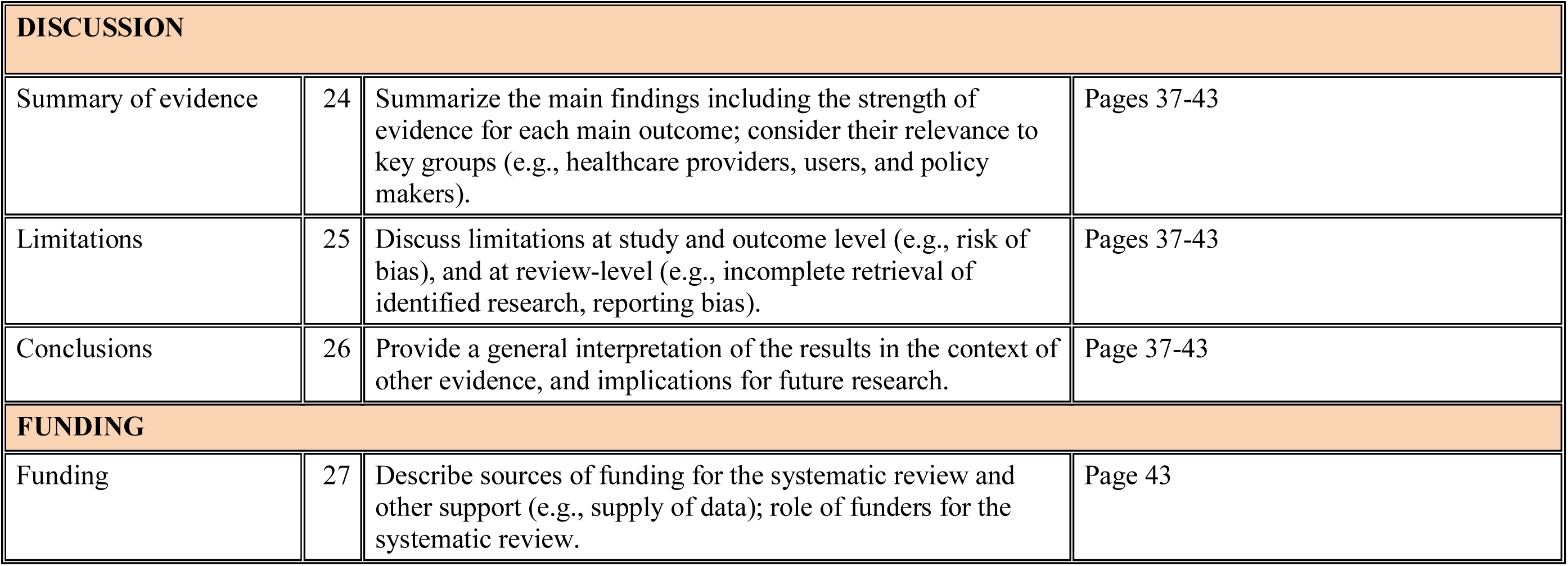

4. PRISMA 2009 Checklist for the Insulin Suppression by Octreotide meta-analysis.

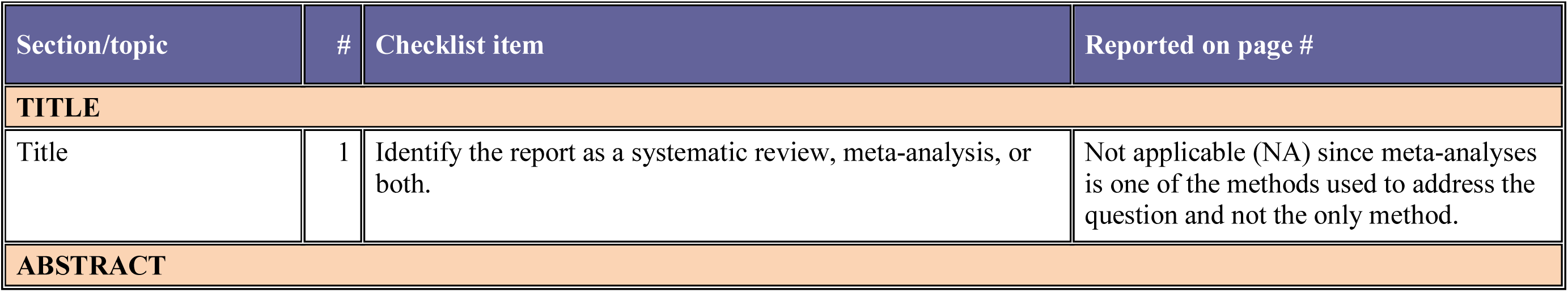
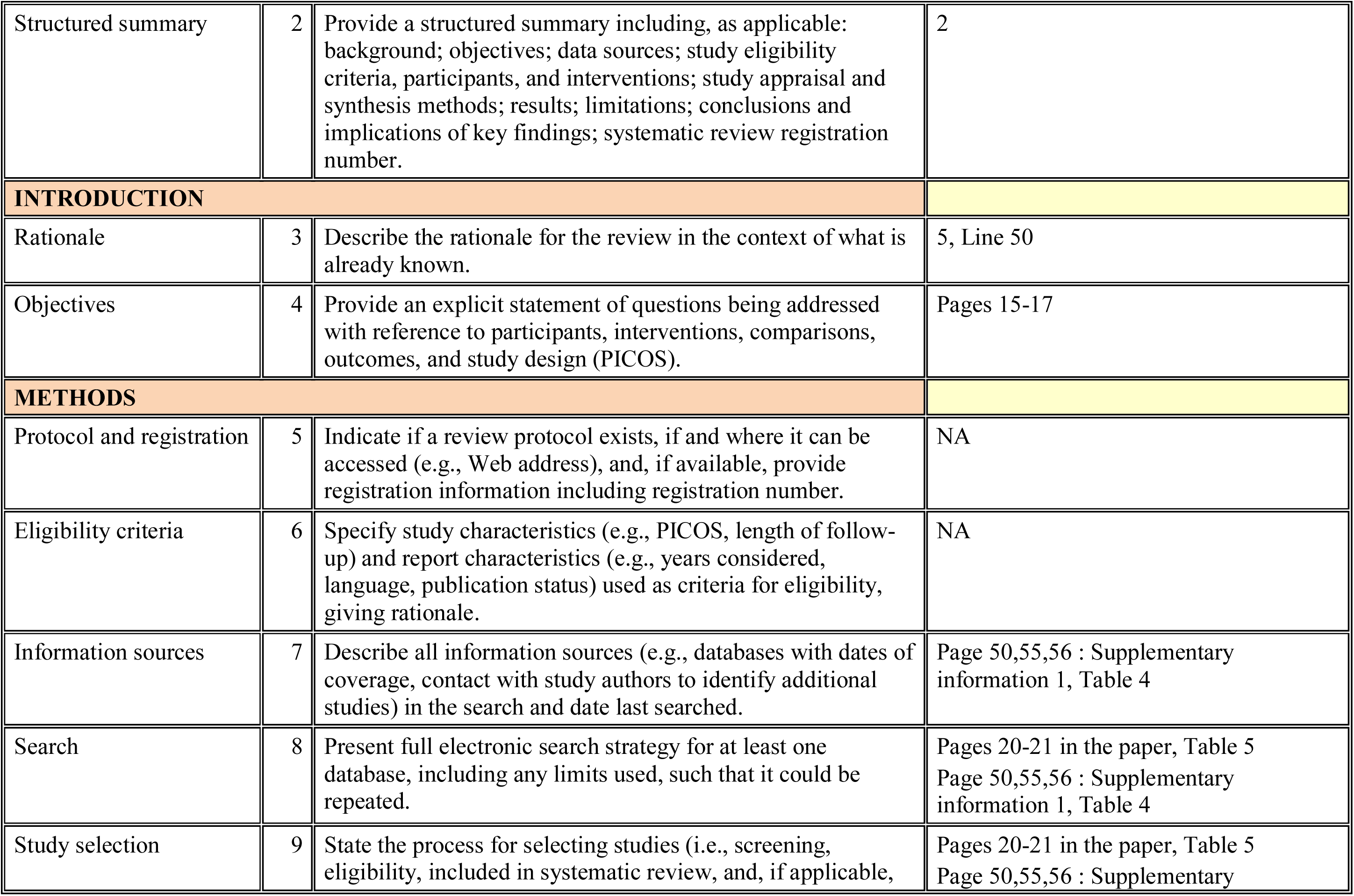
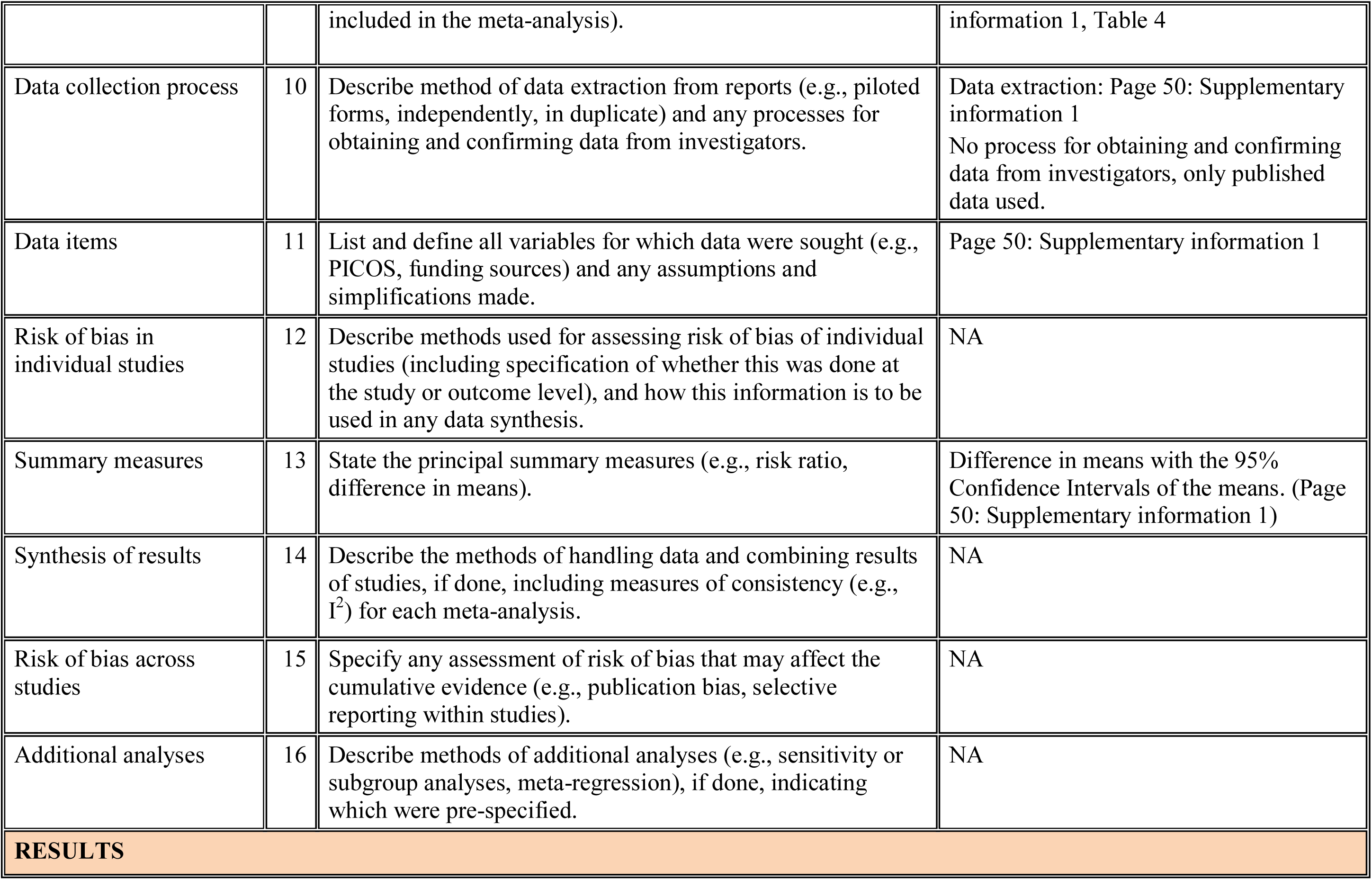
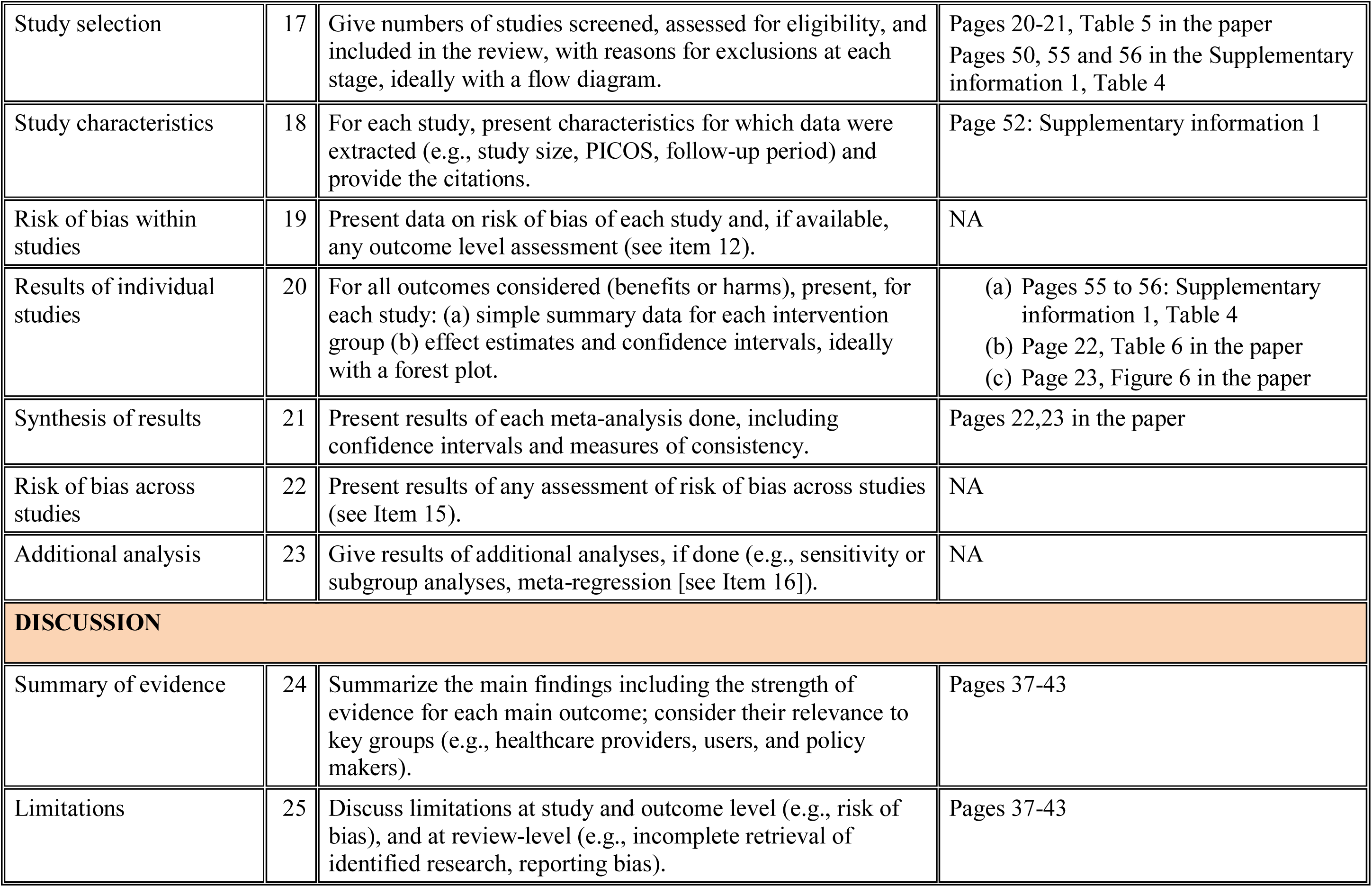
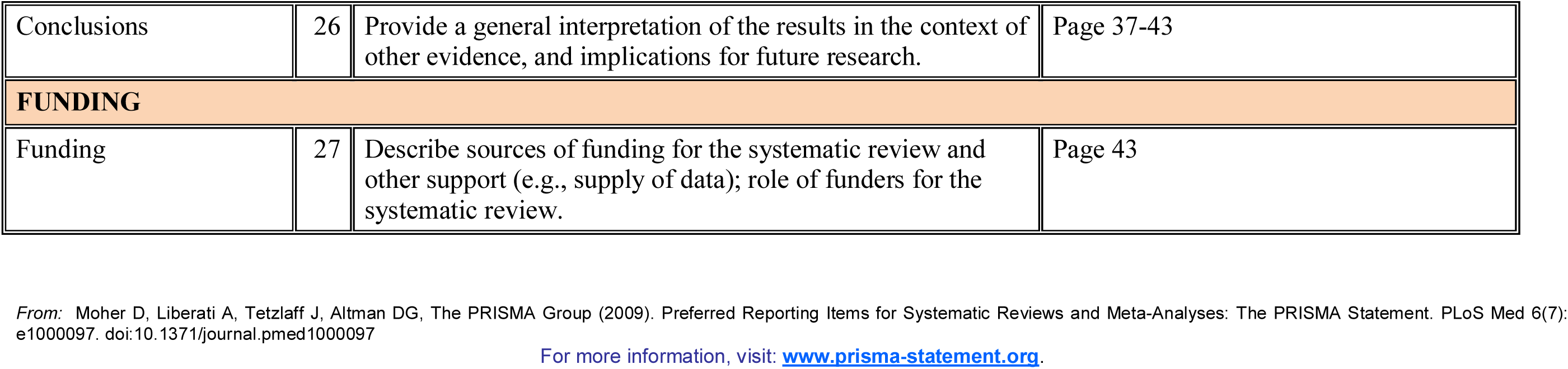

